# Rising Temperature Drives Tipping Points in Mutualistic Networks^†^

**DOI:** 10.1101/2022.04.21.488997

**Authors:** Subhendu Bhandary, Smita Deb, Partha Sharathi Dutta

## Abstract

The effect of climate warming on species physiological parameters, including growth rate, mortality rate, and handling time, is well established from empirical data. However, with an alarming rise in global temperature more than ever, predicting the interactive influence of these changes on mutualistic communities remains uncertain. Using 139 real plant-pollinator networks sampled across the globe and a modelling approach, we study the impact of species’ individual thermal responses on mutualistic communities. We show that at low mutualistic strength plant-pollinator networks are at potential risk of rapid transitions at higher temperatures. Evidently, generalist species plays a critical role in guiding tipping points in mutualistic networks. Further, we derive stability criteria for the networks in a range of temperatures using a two-dimensional reduced model. We identify network structures that can ascertain the delay of a community collapse. Until the end of this century, many real mutualistic networks can be under the threat of sudden collapse, and we frame strategies to mitigate them. Together, our results indicate that knowing individual species thermal responses and network structure can improve predictions for communities facing rapid transitions.

## Introduction

The rate of increase in global temperature over the past 25 years is approximately four times greater than the rate of increase over the past 150 years as a whole (Post 2013). An alarming rise in global temperature is one of the major aftermaths of human influence on climate that disrupts widespread population dynamics (Parmesan and Yohe 2003, Fussmann et al 2014, Malhi et al 2020, Trisos et al 2020). The consequences of such disruptions on species abundances, interactions, and community collapse are poorly understood (Yasuhara et al 2008, Soultan et al 2019, Yeakel et al 2014, Dakos and Bascompte 2014). Predicting species responses to ongoing global warming is of sheer importance for the management and conservation of ecosystems. Until now, little is known as to how increasing temperature influences species dynamics in complex communities (Amarasekare 2015, Vasseur and McCann 2005, Van de Wolfshaar et al 2008). A few recent studies exist highlighting the effects of climate warming on food webs (Yacine et al 2021, Bonnaffé et al 2021). These studies confirm the complex changes along trophic levels caused by warming, and eco-evolutionary feedbacks as a subsequent conservation policy to preserve biodiversity. Detecting the response of communities encompassing species of different genus connected in terms of cooperation and competition remains largely elusive (Bastolla et al 2009, Sugihara and Ye 2009, Stone 2020, Brose et al 2005). Specifically, recognizing the effects of warming on the structure and function of mutualistic communities is crucial (Binzer et al 2016, Takemoto and Kajihara 2016, Albrecht et al 2018, Nagaishi and Takemoto 2018, Bascompte et al 2019).

Mutualism is the ecological interactions between different species belonging to two distinct taxa cooperating for mutual welfare and services (Bascompte and Jordano 2013, Díaz-Castelazo et al 2010, Gómez et al 2011, Domínguez-García and Munoz 2015). Mutualism has been principal to the survival of several species and an essential component of biogeochemical cycles, such as carbon and major nutrient cycles. Mutualistic interactions are also primary to plant colonization and evolution of eukaryotic cell (Potts et al 2010, Wilson et al 2009). As mutualism binds a multitude of species to a common fate, benefiting them, they also hold these species at potential risk of extinction on being exposed to a degrading environment (Ferrière et al 2004). Species in mutualistic networks, ranging from manufacturer-contractor to plant-pollinator networks, interact for enhanced productivity. Although mutualistic networks being widely heterogeneous in terms of interactions per species, these networks possess a well-defined connectivity distribution and structural pattern (Bascompte and Jordano 2013, Valdovinos 2019). Plant-pollinator networks usually share a higher degree of nestedness for a given connectance (Bascompte et al 2003, Astegiano et al 2015). These structural properties have implications on the robustness of a mutualistic community (Revilla et al 2015, Fortuna and Bascompte 2006, Nagaishi and Takemoto 2018). Loss of resilience of mutualistic networks to extinction threats has been observed previously in the face of climate warming (Nagaishi and Takemoto 2018, Bascompte et al 2019). Recent studies have reported that nestedness, which ensures a cohesive core and asymmetric degree distribution, is mainly responsible for the stability and of mutualistic networks (Mariani et al 2019). In all probability, mutualistic networks have an optimal structure that maximizes ecosystem productivity and ensures network stability in deteriorating environmental conditions (Takemoto and Kajihara 2016). Due to the current global warming, the variation in the habitat temperature and a network’s structural properties that influence the proximity of a critical transition demands a careful investigation.

Critical transition or tipping in an ecosystem is characterized by sudden, large, often irreversible, and unexpected shifts from a steady-state to another alternative steady-state due to a parameter drifting (Dai et al 2012, Veraart et al 2012, Drake and Griffen 2010, Carpenter et al 2011). Whilst critical transitions are relatively easy to forecast when a leading species or a small number of species determine the state of an ecosystem, this is not the case for complex communities where interactions between many species determine ecosystems dynamics. Critical transitions can occur in mutualistic communities due to the positive feedback between mutually beneficial species; in plant-pollinator communities, a decline in pollinator abundances can negatively affect plant abundances, which in turn is bad for pollinators (Dakos and Bascompte 2014, Tylianakis and Coux 2014, Lever et al 2020). Mutualistic networks being an integral component of the ecosystem (Bascompte and Jordano 2007); predicting tipping points in mutualistic networks due to increasing mean habitat temperature is critical (Lever et al 2014). This will also aid in recognizing the vulnerable species in a network and have widespread management implications. It remains a challenge to detect tipping points and develop mitigation strategies to foster an early recovery of mutualist species, while model parameters and process rates are functions of temperature.

Considering species’ biological rates and parameters as constants neglects the effect of rising temperature on network stability, ignoring which we may miss out on information crucial to fore a community collapse. To our knowledge, no theoretical/modeling studies have yet considered the effects of species individual thermal responses on communities exhibiting mutualism. It remains unclear how species dynamics on being exposed to high-temperature conditions can trigger cascades of extinction, thereby a community collapse in a mutualistic network (Burkle et al 2013, Six 2009). Here, we investigate sudden transitions in 139 real-world mutualistic networks (from the Web of Life: Ecological Networks Database “http://www.web-of-life.es/”) subject to varying degrees of temperature. We develop a mutualistic network model incorporating species individual thermal responses. This is particularly important as mutualistic interactions between plant-pollinator communities involve plant-visiting ectothermic insects that are sensitive to temperature variations (Shrestha et al 2018).

Here, we show the appearance of tipping points at high temperatures in a plant-pollinator network, where by a network’s state abruptly shifts to an alternate state as the driver of pollination declines. We focus on the correlation between climate warming and hysteresis, irrespective of the complexities in a higher dimensional system. Encouragingly, we found that mutualistic plant-pollinator networks with an optimal structural property can withstand harsh warming conditions and exhibit increased resilience to perturbations. However, the required optimal structure for sustenance varies with the level of environmental deterioration. We also consider the effect of node loss or forbidden links due to critically low abundance or functional dissimilarity between species exhibiting mutualism. Further, we determine the role of the generalist species in triggering a community collapse. Preventing loss of such species has the potential to prevent or delay a community collapse at high temperatures. Stability analysis is imperative to explore the functioning of an ecological system, but this is theoretically challenging for a higher dimensional nonlinear system. We perform stability analyses of the reduced two-dimensional model using dominant eigenvalue of the Jacobian matrix evaluated at the steady states, which allowed us to understand the dynamics of the higher dimensional network with temperature variation (Jiang et al 2018). Overall, our findings underline that efforts to mitigate climate warming and suitable conservation policies can manage the extinction risk of mutualistic communities.

## Models and Methods

We perform our analysis on real plant-pollinator networks differing in their structural properties (e.g., connectance, nestedness), dimensionality, and species variety. These networks are also diverse depending on their geographical location and climatic zones. We employ the interaction matrices for these networks into a mutualistic network model. Existing mutualistic network models, often described by a set of first-order differential equations, represent the dynamics of plant-pollinator communities (Lever et al 2014). However, they do not include species biological rates and parameters as temperature-dependent functions. Here, we develop a network model incorporating the influence of temperature on species biological rates and parameters (i.e., growth rate, mortality rate, and handling time). We consider *A_i_* and *P_i_* as the pollinator and plant abundances in the *i*-th node of a network, respectively. The model representing a group of *S_A_* pollinator species and *S_P_* plant species has the following form:

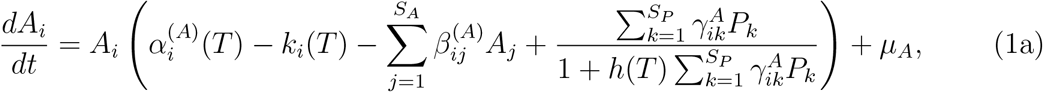

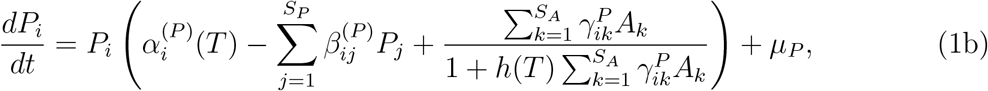

where 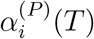 and 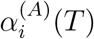 represent the temperature-dependent intrinsic growth rate of i-th plant and pollinator, respectively. *k_i_*(*T*) denotes the decay rate of pollinators, *h*(*T*) is the handling time, and *β_ij_* represents the competition strength between species. *μ_P_* and *μ_A_* represent the immigration factor of plants and pollinators, respectively, and are incorporated in order to prevent underflow errors or allow re-establishment of otherwise extinct species. However, these terms do not influence the qualitative dynamics of the system (Jiang et al 2018, 2019, Lever et al 2014, Meng et al 2020). Mutualistic interaction may be present or absent, and its strength is denoted by 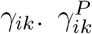 and 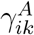 represent the strength of mutualistic interaction of plant and pollinator, respectively. *γ_ik_* is a function of the nodal degree *d_i_* and takes the following form:

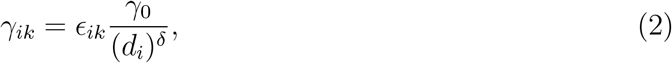

where *ϵ_ik_* represents the entries of the network interaction matrix (obtained from 139 real-world networks); *ϵ_ik_* = 1 if a link is present between the *i*-th plant and *k*-th pollinator species, and is 0 otherwise. *γ*_0_ is the average mutualistic strength, and *δ* modulates the trade-off between the interaction strength and the number of interactions. No trade-off (i.e., *δ* = 0) is the case of mutualistic interaction strengths not influenced by the network structure. In contrast, a full trade-off (*δ* = 1) assumes that benefits attained by species from mutualism are independent of the network topology. In real scenarios, one often assumes a moderate *δ* value, and here we consider *δ* = 0.5 for simplicity. 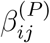 and 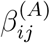 represents the competition strength between *i*-th and *j*-th plant and pollinator, respectively which behave synergistically with temperature beyond optimum (Amarasekare and Coutinho 2014). We consider the intraspecific competition *β_ii_* = 1 and interspecific competition *β_ij_*(*i* ≠ *j*) = 0 (however *β_ij_* can take any value in (0,1), i.e., intraspecific competition higher than interspecific competition).

### Dependence of species process rates and parameters on temperature

Based on empirical evidence, recent studied have confirmed species biological rates (e.g., birth rate, death rateand parameters (e.g., handling time) as functions of temperature (Scranton and Amarasekare 2017, Uszko et al 2017, Kaur and Dutta 2020). Here, we consider temperature dependent species growth rate *α_i_*(*T*) exhibiting a unimodal symmetric response represented by a Gaussian function (Scranton and Amarasekare 2017, Hatfield and Prueger 2015):

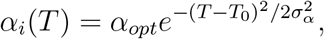

where *T*_0_ is the temperature at which the value of *α_i_*(*T*) is optimal and equals *α_opt_*. *σ_α_* denotes the performance breadth. For simplicity, the intrinsic growth rate of the plant and pollinator species are considered equal (Rohr et al 2014, Lever et al 2014). The handling time *h*(*T*) of the pollinators obeying Holling type-II functional response exhibiting a hump or a U-shaped relationship with temperature can be represented by a Gaussian function (Uszko et al 2017):

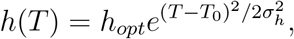

where *h_opt_* represents the value of *h*(*T*) at the optimum temperature *T*_0_. *σ_h_* denotes the performance breadth. The per capita mortality rate of pollinators *k_i_*(*T*) are observed to follow the Boltzman-Arrhenius relationship (Scranton and Amarasekare 2017), and formulated as:

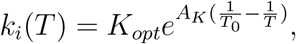

where *K_opt_* is the mortality rate at the optimum temperature *T*_0_ and *A_K_* is the Arrhenius constant. Reduced pollinator population or an increase in pollinator mortality due to environmental factors is modulated by the term *k_i_*. The functional response curves for the species birth rate, handling time, and death rate are plotted in *SI Appendix*, Section S1, Fig. S1.1)

We study dynamics (collapse and the possibility of subsequent recovery) of the network model (1) when birth rate, death rate and handling time (see *SI Appendix*, Section S1, Fig. S1.1) are exposed to rising temperature in the range 0 – 40°*C*, for a fixed average mutualistic strength *γ*_0_. We find the critical temperature *γ*_0_ value for a specific *γ*_0_ at which the system undergoes an abrupt transition to an extinction state and thus estimating species tolerance at different *γ*_0_. By varying temperature in the range 0 – 40^0^*C* in the forward (increasing) as well as backward (decreasing) direction, we examine the point of collapse and recovery, respectively. The difference in the point of collapse and recovery provides us with a metric to study the hysteresis among plant and pollinatorpopulations. To understand the role of network structural properties on the plant-pollinator dynamics, we calculate nestedness using the NODF (Nestedness metric based on overlap and decreasing fill) measure (Lever et al 2014), modularity (Fortuna et al 2010), and connectance(Box 1). All parameters, including temperature range and variability in *γ*_0_ are chosen from the previous studies (Lever et al 2014, Jiang et al 2019). The system is solved numerically using the fourth-order Runge-Kutta method with adaptive step size, and the model is run till a stationary state has reached.

#### Box 1. Glossary

##### Connectance (*C*)

It is the ratio of realized interactions to all possible interactions, 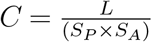, where *L* is the number of realised interactions (links connecting plants and pollinators), *S_P_* and *S_A_* denote the number of plants and pollinators, respectively.

##### Nestedness (*N*)

A bipartite network (usually represented as a matrix) is said to be nested when components having a few items in them (locations with few species, species with few interactions) have a subset of the items of components with more items. In other words, nestedness describes the extent to which interactions form ordered subsets of each other. We calculate nestedness using the Nestedness metric based on Overlap and Decreasing Fill” (NODF) measure, defined as:

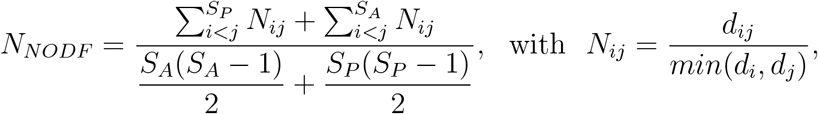

where *N_ij_* is the nestedness of species pair *i* and *j*. *d_i_*, *d_j_* are the number of ones in row *i* and *j*, and *d_ij_* is the number of shared interactions between rows *i* and *j* (so-called paired overlap). Ecologically *d_ij_* is the number of times species *i* and *j* interact with the same mutualistic partner.

##### Modularity (*Q*)

It is the degree to which densely connected compartments within a network can be decoupled into separate clusters interacting more within themselves compared to interaction across clusters.

For a bipartite network represented by the matrix *B*, the modularity metric is expressed as:

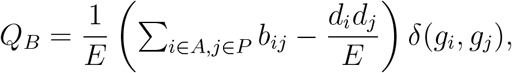

where *b_ij_* is the element in *B* representing a link (i.e., *b_ij_* = 1) or no link (i.e., *b_ij_* = 0) between nodes *i* and *j*, *g_i_* is the module that node *i* belongs to under a certain partition, *d_i_* is the degree of node *i*, *δ* denotes Kronecker’s delta, and *E* is the number of links in the network.

##### Nodal degree

Degree of a node. It is defined as the total number of relationships involving that node.

##### Tipping point

A threshold value at which a dynamical system abruptly shifts from one stable state to another alternative stable state, in response to small stochastic perturbations.

### Dimension reduction of the network model

In a higher dimensional network model having multiple variables, finding the analytical expressions for equilibrium points and determining their stability might be cumbersome. Hence, for mathematical analysis of the mutualistic network model (1), we reduce it in two-dimension by considering averaged value of mutualistic interaction strength among plants and pollinators (Tu et al 2021, Jiang et al 2018). The dimension reduced model can be written as:

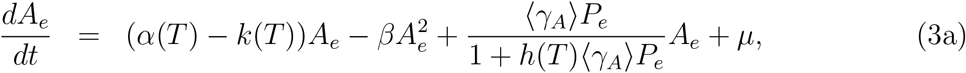

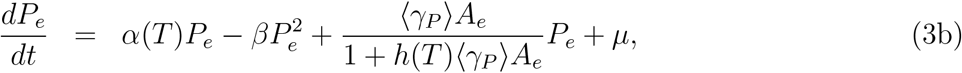

where *A_e_* and *P_e_* are the effective abundances of pollinators and plants, respectively; *α*(*T*) represents the temperature-dependent effective growth rate for the network, *h*(*T*) denotes the effective handling time, *β* represents the combined effect of intraspecific and interspecific competition, 〈*γ_A_*〉 and 〈*γ_P_*〉 denote the effective mutualistic strength of pollinators and plants, respectively, *k*(*T*) is the species death rate in averaging sense and migration effects for the species is represented by *μ* (Jiang et al 2018) (for further details, see *SI Appendix*, Section S2). The dimension reduction technique used here is limited by the inability to incorporate heterogeneity at the nodes (in terms of parameters). However, it enables the study of nonlinear phenomena; such as bifurcations (Ott 2002), basin structures (Grebogi et al 1983) and transient chaos (Hastings and Higgins 1994, Hastings 2001), etc., which are otherwise impossible in the higher dimensional network. As shown in *SI Appendix*, Section S2, Fig. S2.1, the reduced system well depicts the qualitative dynamics of the model, including the dynamics near a tipping point.

## Results

### Temperature driven sudden transitions in mutualistic networks

We demonstrate the role of temperature in driving tippings in mutualistic networks of varied dimensions, sampled across the globe. As depicted in Fig. 1, with gradual change in temperature a network can undergo sudden transition from one stable state to an alternate stable state. However, this result changes with the strength of mutualistic interaction (*γ*_0_). The abundance of pollinators varies for change in mean habitat temperature, and can undergo an abrupt shift beyond optimum. With an increase in *γ*_0_, the threshold temperature at which the system collapses increase considerably. Further increase in *γ*_0_ beyond 1.5 prevents sudden community collapse in the considered feasible temperature range.

**Figure 1.**
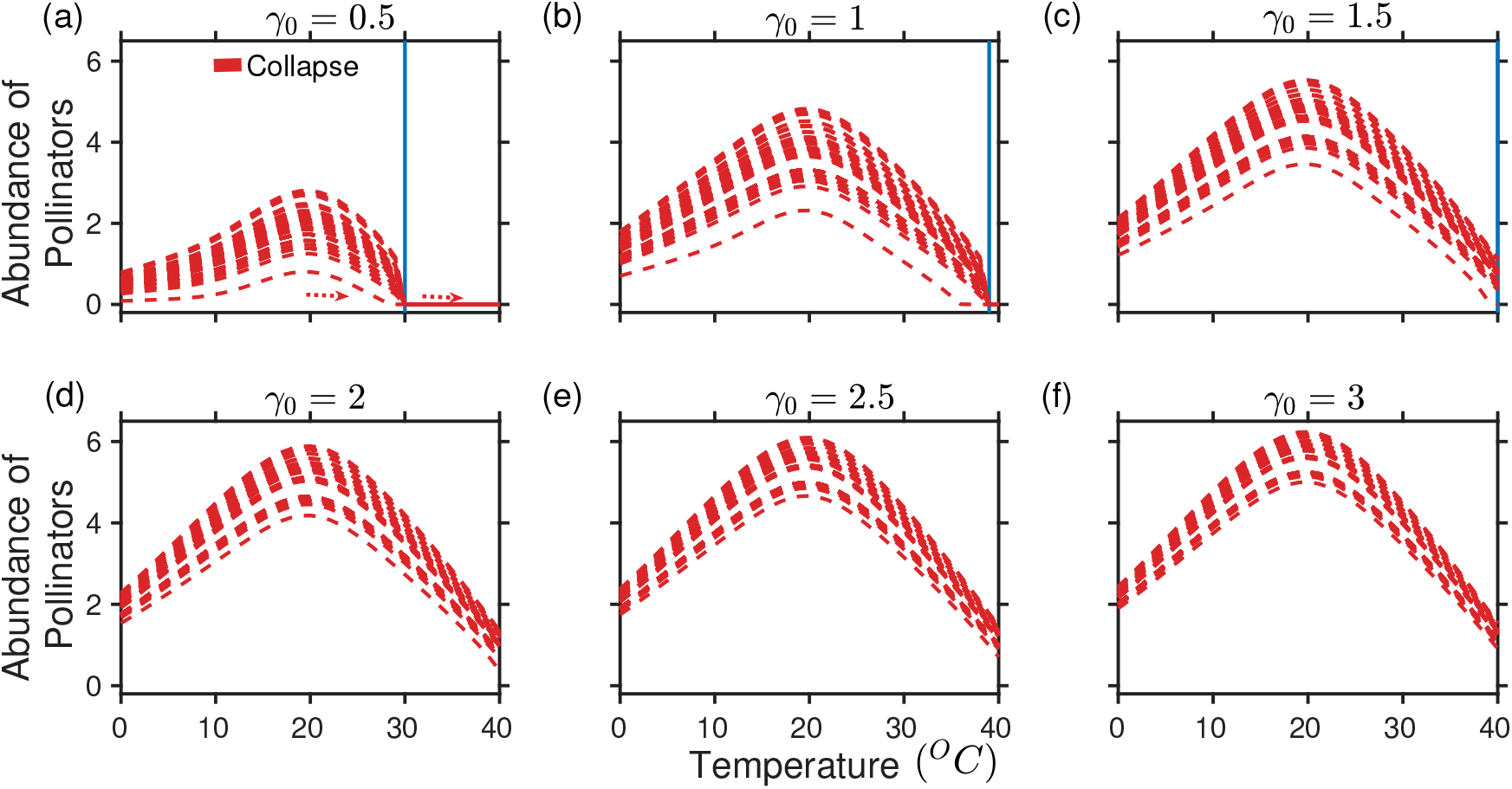
Higher temperature can trigger catastrophic transitions at different interaction strengths (*γ*_0_): (a)-(c) On increasing the temperature in the range (0 – 40°*C*), for *γ*_0_ = 0.5 to *γ*_0_ = 1, the abundance of pollinators encounters catastrophic transitions. (d)-(f) At or beyond *γ*_0_ = 1.5, sudden community collapse is averted, despite there is a gradual drop in population abundance with increasing temperature. The blue vertical line marks the occurrence of critical transition. Each lines in sub-figures represent the abundance of pollinators at each node. The above result is obtained for network *M_PL*_006 with *S_A_* = 61 and *S_P_* = 17 (for details, “http://www.web-of-life.es/”). The taxonomic details of the above network is presented in *SI Appendix*, Section S1, Table. S1.2. The parameter values are 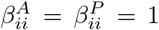, *δ* = 0.5, *μ_A_* = *μ_P_* = 10^−4^, and the other parameters 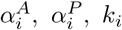 are obtained from their respective response function (unless stated, the network used and these values are same in rest of the figures).

Our findings suggest that high temperatures can trigger critical transitions in a mutualistic community. This can otherwise be debilitated by lowering the degree of warming or prevent/delay a transition by maintaining the requisite *γ*_0_. However, at a fixed *γ*_0_, striving to recover species demands lowering temperature at least by 2 – 3^0^*C* or more which is extremely challenging in the face of rapid global warming. On the otherhand *γ*_0_ may be weak or strong depending upon various global environmental factors (Tylianakis et al 2008, Saavedra et al 2013), although direct relations are unknown. In line with existing literature, we analyze the effects of both way change in interaction strength *γ*_0_ and their impact on network collapse at different temperatures.

As depicted in Fig. 2, for low mean habitat temperature up to optimum, the abundance of pollinators gradually collapse at comparatively lower *γ*_0_ values. Beyond this, the community encounters sudden collapse even at relatively high *γ*_0_ values. Likewise, increasing *γ*_0_ in the range (0 – 3), it is observed that the system recovers early till optimum temperature, beyond which recovery is delayed. In the temperature range 32 – 40°*C*, the system fails to recover for any feasible *γ*_0_ value (0 – 3) and encounters one point collapse. We further observe that the point of community collapse and recovery are the same and differ at and beyond 28°*C*. Clearly, the critical *γ*_0_ value that controls both the collapse and recovery of a system is influenced by the change in temperature (Fig. 2). In order to allow the system to recover, we increase *γ*_0_ in the direction (0 – 3), opposite to that which causes collapse. This leads to the formation of a hysteresis loop (Figs. 2(d)-2(f)). The hysteresis loop is created first at approximately 28°*C*, which was otherwise absent at low temperatures. More explicitly, in Fig. 2(f), at 40°*C* the system collapses at *γ*_0_ = 1.1 and does not recover on improving conditions post-collapse.

**Figure 2.**
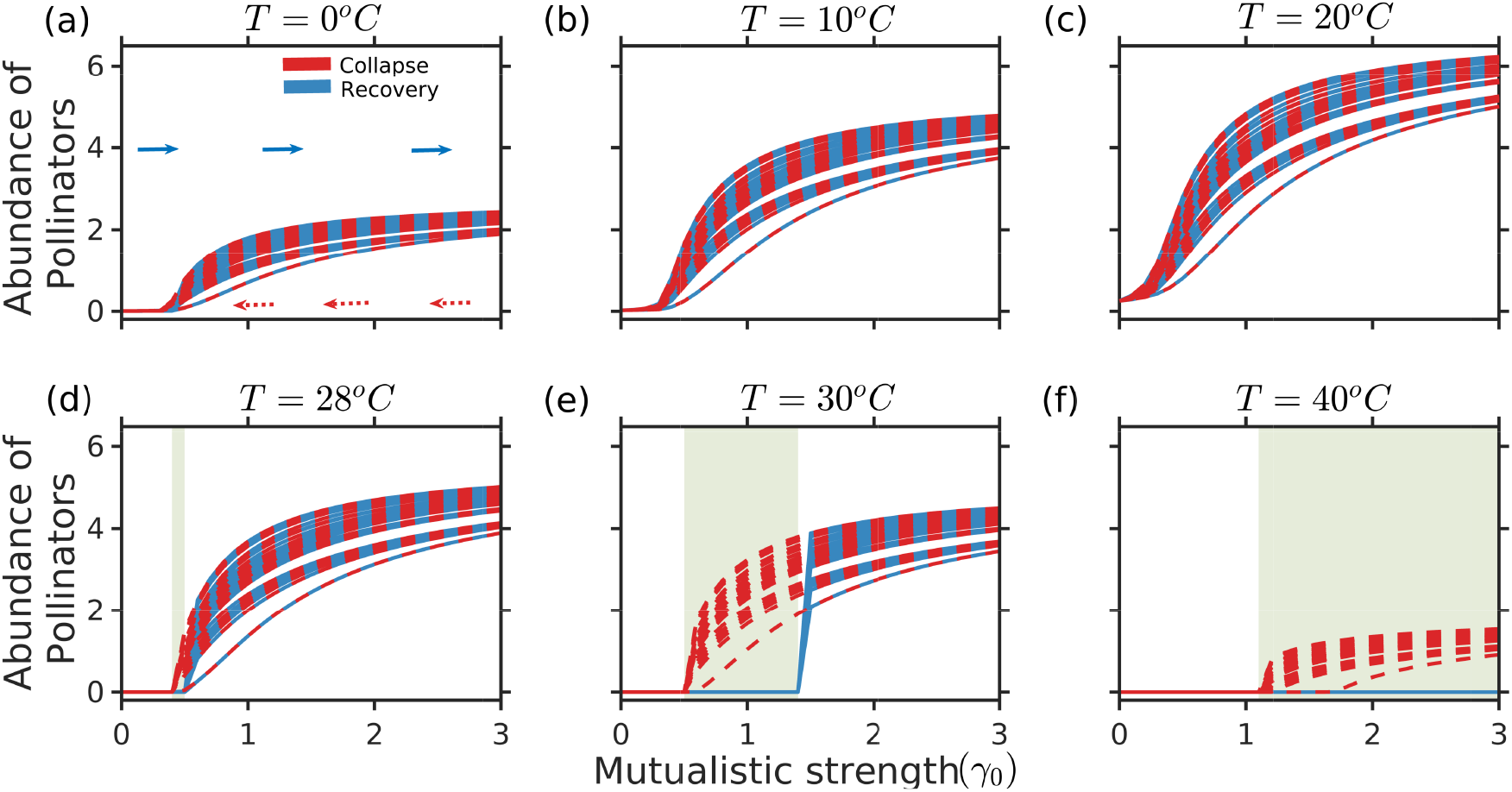
Catastrophic collapse in mutualistic networks for variation in mutualistic strength (*γ*_0_): (a)-(b) On decreasing the mutualistic strength *γ*_0_ in the range (3-0) (red), at low to moderate temperature till 20°*C*, the abundance of pollinators encounters a non-catastrophic transition. On increasing *γ*_0_ in the range (0-3), the system recovers to its previous state. (c) At 20°*C*, community collapse is averted, despite there is a drop in abundance at low *γ*_0_ values. (d)-(f) As the temperature is further increased, a gradual shift transforms into a rapid collapse at a low *γ*_0_ value. As *γ*_0_ increases, striving to recover species (in the forward direction 0 – 3 (in blue)), a hysteresis loop formation is observed, and the width of this loop increases with an increase in warming. At very high temperature (40°*C*) the system does not recover. Results for other networks are presented in *SI Appendix*, Section S1, Fig. S1.2.

### Temperature-structure interplay influences tipping

In this section, we study the combined effect of the structure of a mutualistic network and rising temperature on influencing tipping points. Structural property, viz, nestedness, influences the tipping, which is prevalent at higher temperatures. Higher trade-off (*δ* value) accounts for the effects of asymmetry. However, mixed effects of trade-off are observed at low temperature across networks (*SI Appendix*, Section S1, Fig. S1.9). In Fig. 3, we show results of 139 networks having nestedness varying in the range (0-0.84) allowing the networks to be exposed to mean habitat temperature (0° – 40°*C*) (Vasseur et al 2014, Uszko et al 2017, Kaur and Dutta 2020). We find that nestedness plays a critical role in eluding the first point collapse (the point at which abundance of pollinators in at least one of the nodes falls below 1 × 10^−2^) at high temperature but have had mixed effects on the final point collapse (loss of species abundance at all the nodes). The tipping points for highly nested networks occur at substantially low *γ*_0_ values, whilst collapse for networks with low nestedness occurs despite maintaining high *γ*_0_ between species at extreme temperature. At low-moderate temperature, nestedness does not quite influence the tipping. Figure 3 depicts the relation between *γ*_0_, nestedness, and occurrence of tippings at different temperatures. Both being majorly uncorrelated for the temperature change, but exhibit strong negative correlation beyond optimum (*SI Appendix*, Section S1, Fig. S1.10). The methods for calculating nestedness of networks have discrepancies, and there remains a debate on the calculation of nestedness (Astegiano et al 2015, Fortuna et al 2010, James et al 2012, Staniczenko et al 2013, Thomas et al 2015). Nevertheless, nestedness could be correlated to other network properties such as connectance and network size but for comparing our results with previous studies we adopt the NODF measure (Lever et al 2014). Therefore, we analyze the effect of other structural properties of networks, e.g. connectance and network size on the collapse of mutualistic networks with variation in temperatures. The results presented in *SI Appendix*, Section S1, Fig. S1.7(a) show that mutualistic networks with higher connectance experience delayed first point collapse and no prominent effect on the final point collapse (*SI Appendix*, Section S1, Fig. S1.7(b)); consistent with our findings in Fig. 3 across different nestedness values. Furthermore, we present first point and last point collapse of the 139 plant-pollinator networks and show that no prominent effect is visible with variation in the number of nodes in a network (*SI Appendix*, Section S1, Fig. S1.7(c)-(d)), unlike nestedness and connectance. Also as observed in (*SI Appendix*, Section S1, Fig. S1.8), connectance and nestedness in the considered 139 plant-pollinator networks-the two structural properties are independent of network size. Therefore for the considered empirical data, either nestedness or connectance may be considered as network structural property for further theoretical explorations. In line with the existing literature, our results also suggest that nested networks are most robust to extinction and habitat loss than their random counterparts. Figure 3(c) shows a positive correlation between nestedness and connectance, while Fig. 3(d) shows modularity is negatively correlated to nestedness. Thus, results indicate high nestedness and connectance as structural properties of a mutualistic network, enabling them to withstand high temperatures and delay tipping.

**Figure 3:**
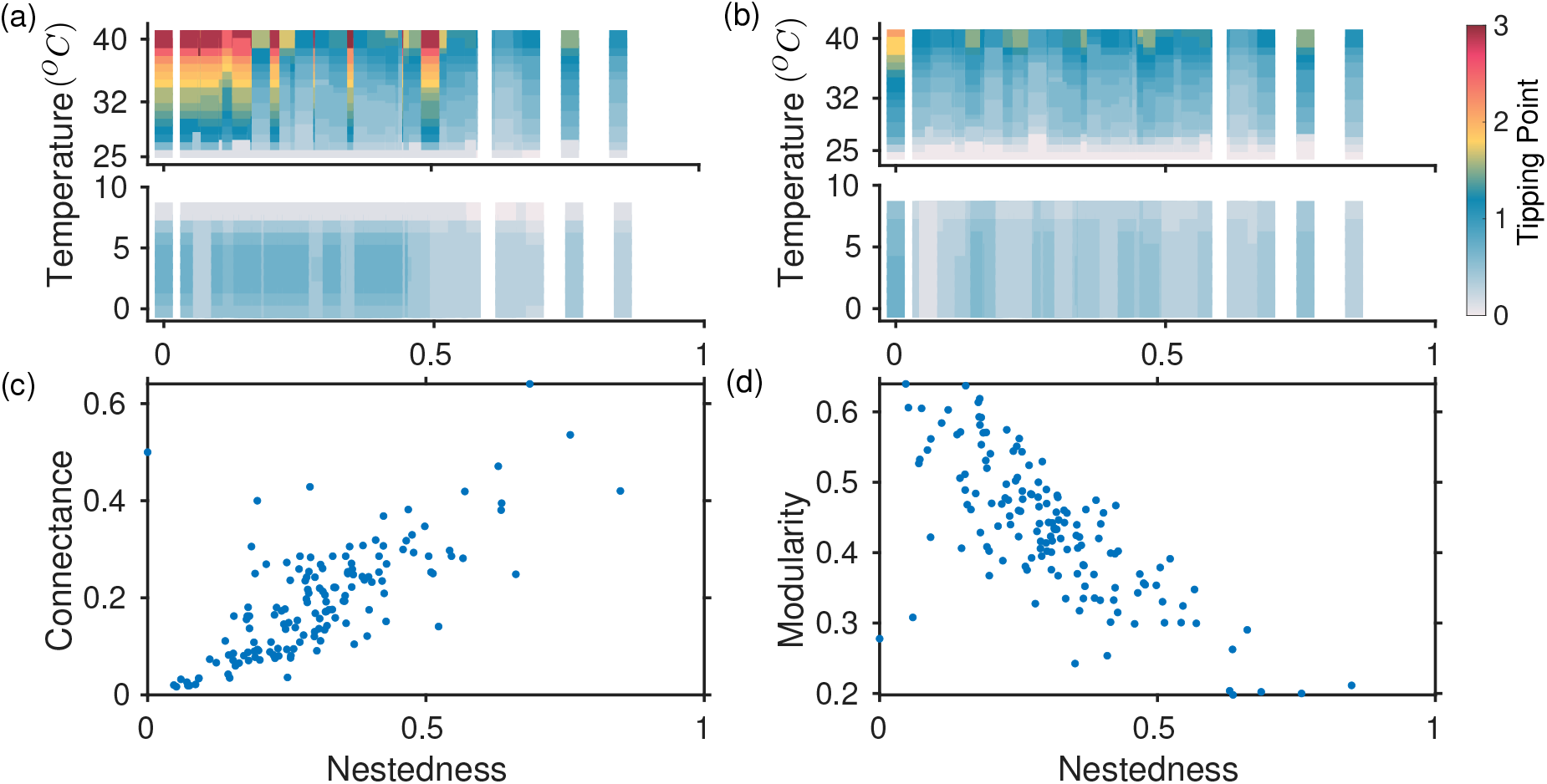
Role of network structure under varied degree of warming in delaying a tipping: (a) First point collapse of pollinators for 139 real plant-pollinator networks. At higher temperatures, nested networks undergo collapse at a considerably lower *γ*_0_ value (which marks the tipping point). (b) Final point collapse of the networks. The colors in the color bar correspond to the *γ*_0_ values in the range (0-3), at which the system undergoes a first (a) and final point collapse (b). Correlation between; (c) connectance and nestedness, and (d) modularity and nestedness of 139 real networks. The results are averaged over 100 independent simulations.

### Effects of node loss and link loss on network dynamics

Here, we investigate if *γ*_0_ affects all species equally or the core of generalists (nodes with higher connectivity) are predominant over their specialist (nodes with lower connectivity) counterparts at higher temperatures. We study network resilience under two types of perturbations. First, we remove fraction *f_P_* of nodes representing plant loss, and second, we remove fraction *f_A_* of pollinators to study link perturbation. In Fig. 4, we study the impact of the more functional plant and pollinator loss, respectively, on a network. In Figs. 4(a)-4(f), we observe that on the removal of a fraction of nodes (plant loss) (Gao et al 2016) in decreasing order of their degree, there is negligible variation in the point of collapse at low to intermediate temperatures. On the contrary, at higher temperature (28°*C* and above) as a larger fraction of generalists (both plants and pollinators) are deleted, the point of collapse of the largest fraction precede that of the smallest by a considerable amount. In Figs. 4(g)-4(l), we study the effect of small link perturbations (fraction of pollinators removed) on the network at various habitat temperatures (0°*C*, 5°*C*, 10°*C*, 30°*C*, 35°*C*, 40°*C*). We observe that the system is resilient to infinitesimal perturbations. However, on further increase in link loss, community collapse is elevated at 40°*C*. Our results indicate that effects of both link loss and node loss are more profound at extreme warming conditions.

**Figure 4.**
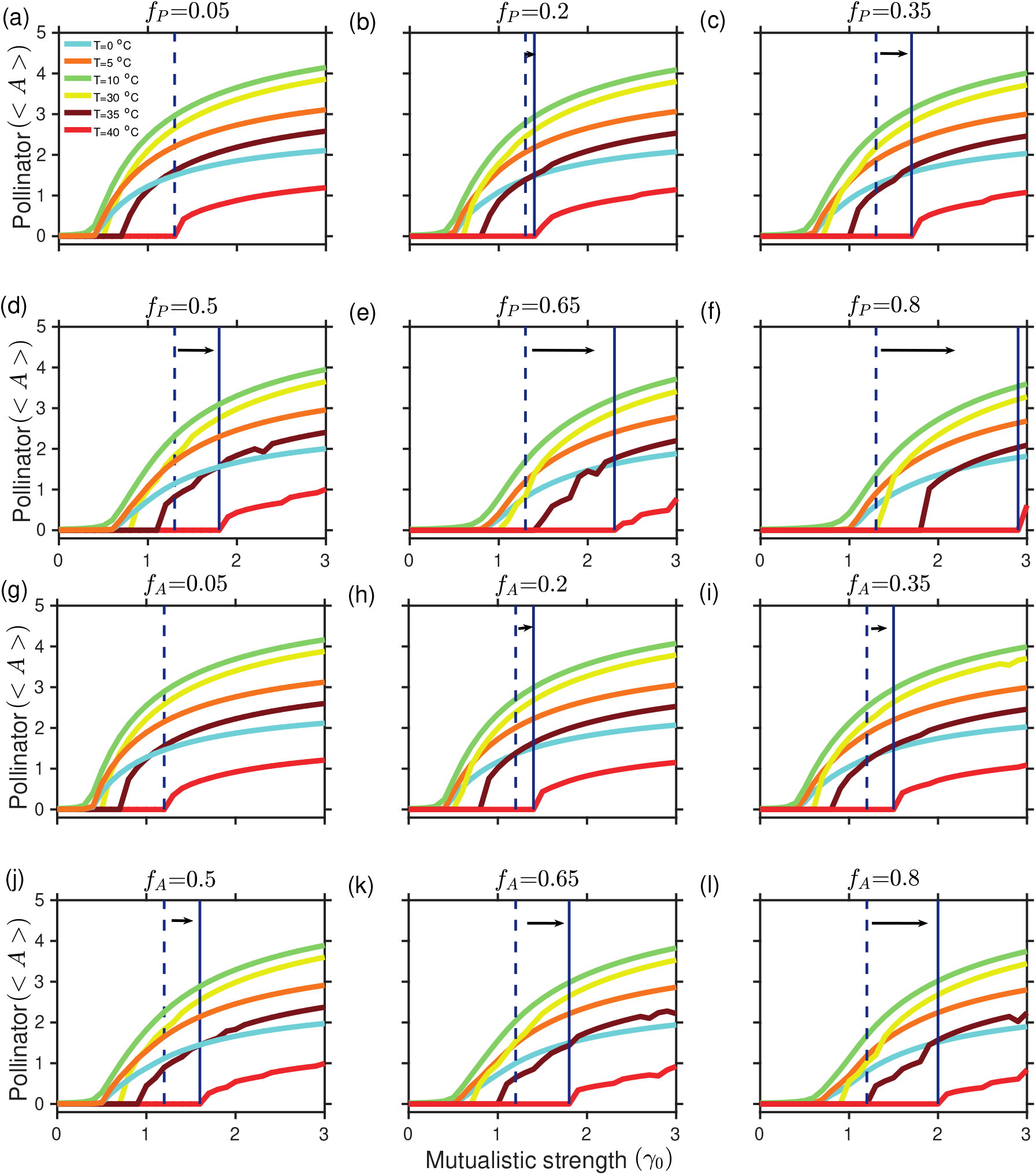
Effects of plant and pollinator loss for varied temperature regimes: The decline in the average abundance of the pollinator community as a fraction of plant (*f_P_*) (a)-(f), and pollinators (*f_A_*) (g)-(l) are removed in decreasing order of their degree at temperatures ranging over the interval 0°*C* – 40°*C* (denoted by different colors in the legend). As *γ*_0_ decreases from 3 – 0, the point of collapse precedes as *f_P_* and *f_A_* are increased further. Plant loss has a more prominent effect on the collapse of the pollinator community as indicated by the distance between dashed and solid vertical lines for respective *f_P_* and *f_A_* values. A large difference is observed in tipping at high temperature (40°*C*) for increase in *f_P_* = 0.8 and *f_A_* = 0.8. The dashed lines in (a)-(l) represent the point of collapse for *f_P_* = *f_A_* = 0.05 at 40^0^*C*. The solid lines in (b)-(f) and (h)-(l) represent the point of collapse for different values of *f_P_* and *f_A_* as mentioned on top of each sub-figures at 40^0^*C*. < *A* > denotes the average abundance of pollinator species.

### Stability analysis of the network model via dimension reduction

In this section, after reducing the temperature-dependent mutualistic network model into a two-dimensional system, we study its dynamics. Mutualistic networks are composed of a large number of species leading to a high phase-space dimension. Analysis of this higher-dimensional model requires insight into the underlying dynamics. This requires incorporating the necessary interactions by applying the weighted approach of obtaining the ensemble average value of *γ*_0_ (*SI Appendix*, Section S2).

Figure 5 presents the average abundance of the pollinator species of the stable steady-state (*SSS*) and unstable steady-state (*USS*), respectively. We observe that the non-trivial stable steady-state extends over the entire temperature for *γ*_0_ varying in the range (0.5 – 3). However, the system is unstable for very high (28° – 40°*C*) and low (0° – 8°*C*) temperatures. The two-dimensional projection reveals that the abundance corresponding to the stable branch is obtained for the maximum range of considered *γ*_0_ in a closed neighborhood of the optimum temperature (Fig. 5). At higher and lower temperatures, *γ*_0_ region corresponding to the *SSS* is reduced compared to the intermediate temperatures, and the area is decreased further as we move away from the optimum temperature. It is evident that species biological responses to temperature is a key factor that governs the stability of mutualistic networks. Whilst increased *γ*_0_ appears to hold back the system in the stable regime until a threshold temperature is reached, the system loses its stability whenever *h*(*T*) > *α*(*T*) and *k*(*T*) > *h*(*T*). Thus, the persistence of species is more favorable when the species growth rate is much higher than the resource limitation rate and handling time is more than the decay rate, indirectly being controlled by the degree of warming.

**Figure 5.**
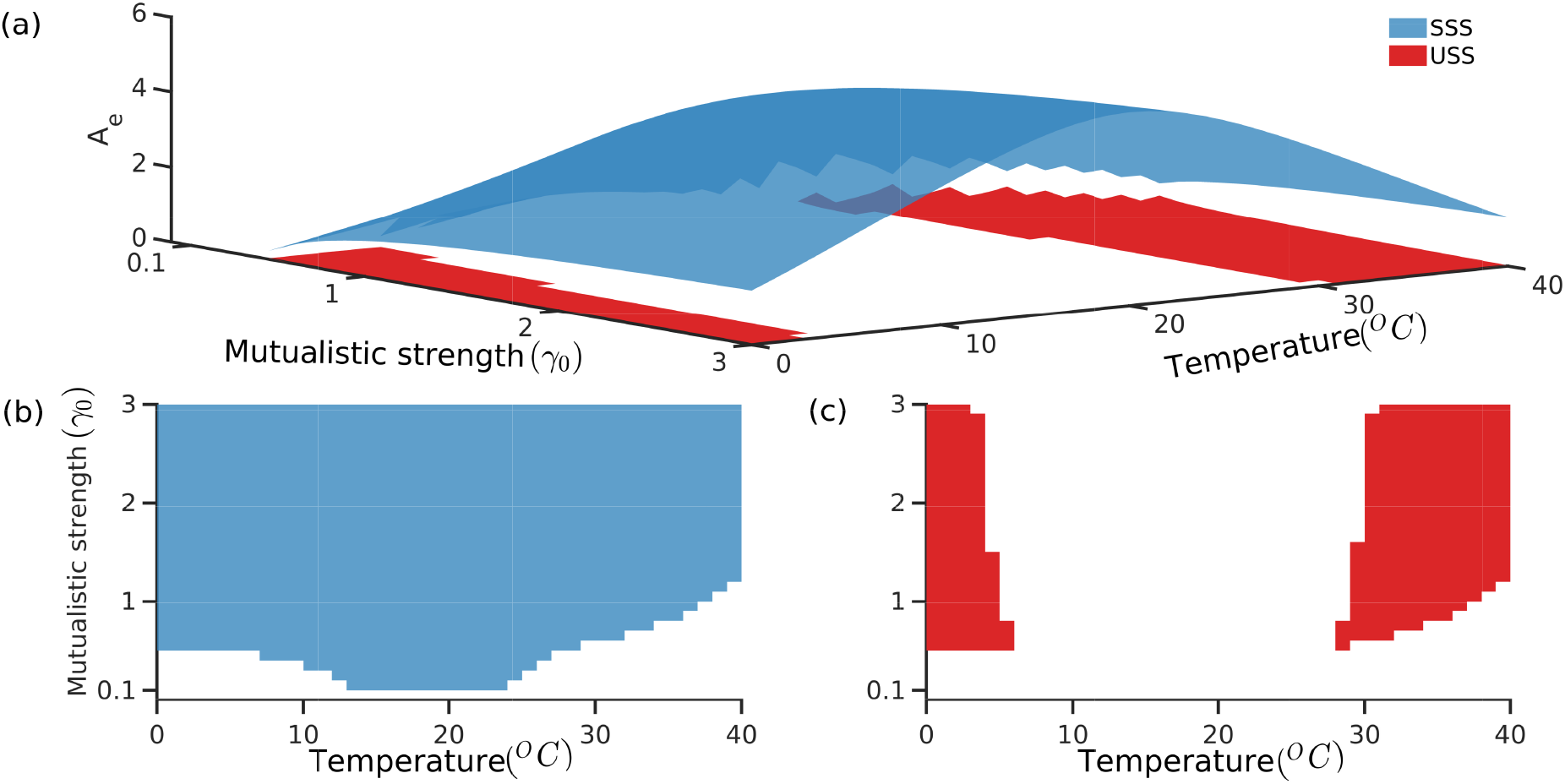
Stable and unstable steady states of the reduced model: (a) shows the stable (blue) and unstable (peach) steady-state surfaces obtained from the reduced model. The ensemble pollinator abundance is plotted as a function of *γ*_0_ in the temperature range 0° – 40°*C*. The 2D projections of the above stable and unstable surfaces depicting the stable and unstable regions at different temperatures are shown in (b)-(c).

Distinctly around the optimum temperature, the system is more stable, as indicated by Fig. 6. Eigenvalues are more negative corresponding to the Jacobian of the *SSS* around the optimum temperature for all *γ*_0_ (*SI Appendix*, Section S2, Fig. S2.2). The behavior remains consistent across all the 139 networks with varying structural properties (Fig. 6). Whilst an increase in *γ*_0_ leads to a more negative dominant eigenvalue, the effects cannot be isolated to the optimum temperature alone and are minimal beyond *γ*_0_ = 2. However, at lower *γ*_0_ (*γ*_0_ = 0.5), no trends are observed. Our results suggest that high mutualistic strength is a mean to maintain network stability at higher temperatures.

**Figure 6.**
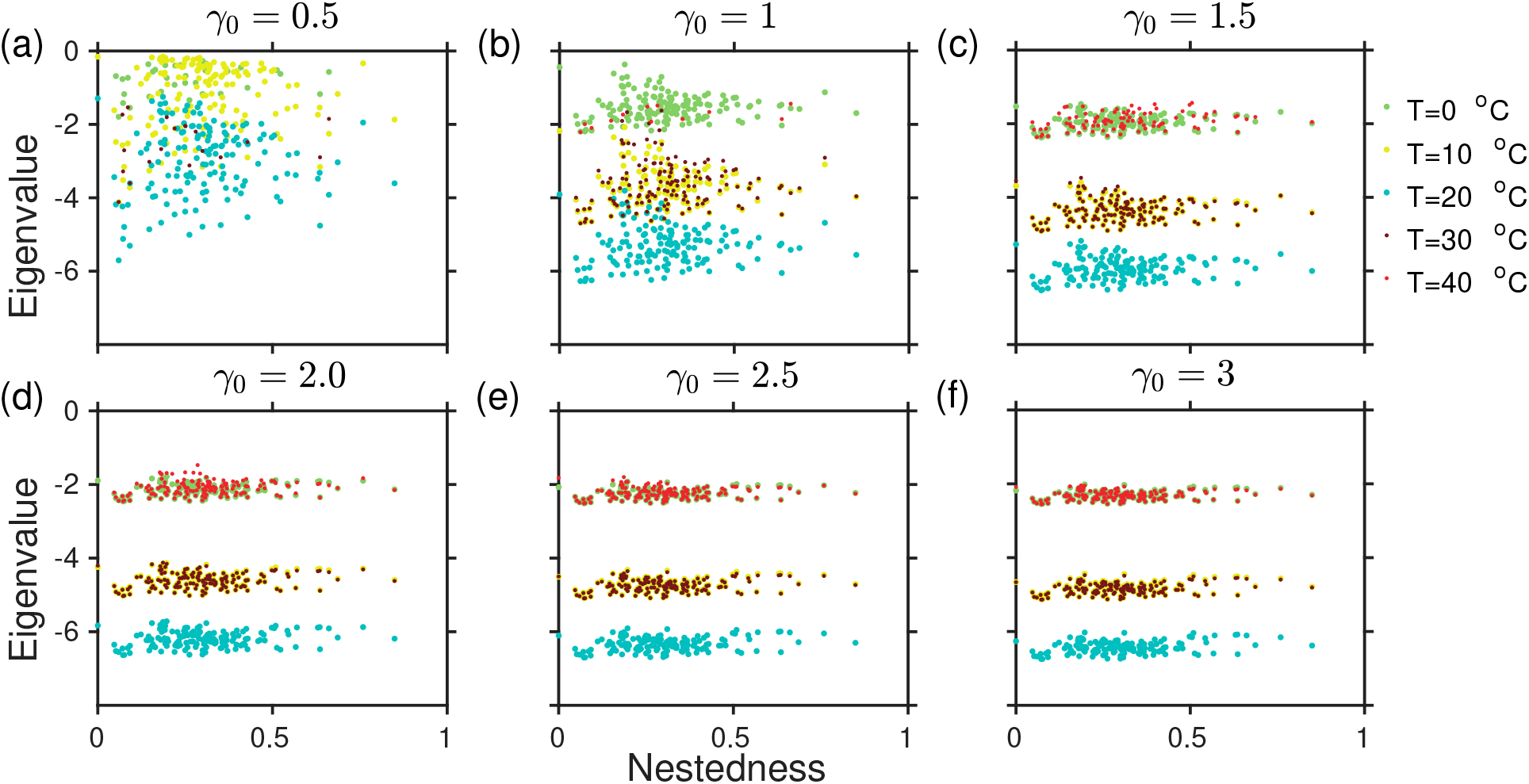
Effect of temperature and network structure on the eigenvalue of the stable steady state: (a)-(f) The eigenvalue of the Jacobian matrix corresponding to the non-trivial steady-state plotted against the nestedness value for all the 139 plant-pollinator networks for a fixed *γ*_0_ mentioned above each panel. Each dot represents a network with the color denoting the respective temperature. The system is always more stable at optimum; stability at higher temperature increases by increasing *γ*_0_, although the critical eigenvalue saturates at *γ*_0_ = 2 and does not increase further.

### Management policies for species sustainability at higher temperatures

We present on the globe (see Fig. 7) regions experiencing temperature in the range (9 – 25°*C*) for the socio-economic pathway (SSP370, CMIP6) (https://esgf-node.llnl.gov/search/cmip6) for the year 2099. This socio-economic pathway represents the scenario where environmental concerns are given low international priority leading to degrading environmental conditions. We plot all the 139 real plant-pollinator networks used for this study on the globe at their respective locations to find out the networks which are at risk of extinction. Only 37 out of 139 networks located in regions experiencing optimum temperatures do not undergo tipping, while the rest 102 networks are at a risk of tipping by the end of 2099 along socio-economic pathway SSP370 (Fig. 7). Although susceptibility of such networks may vary depending on their structure (Fig. 3); they are likely to undergo tipping at different *γ*_0_ values.

**Figure 7.**
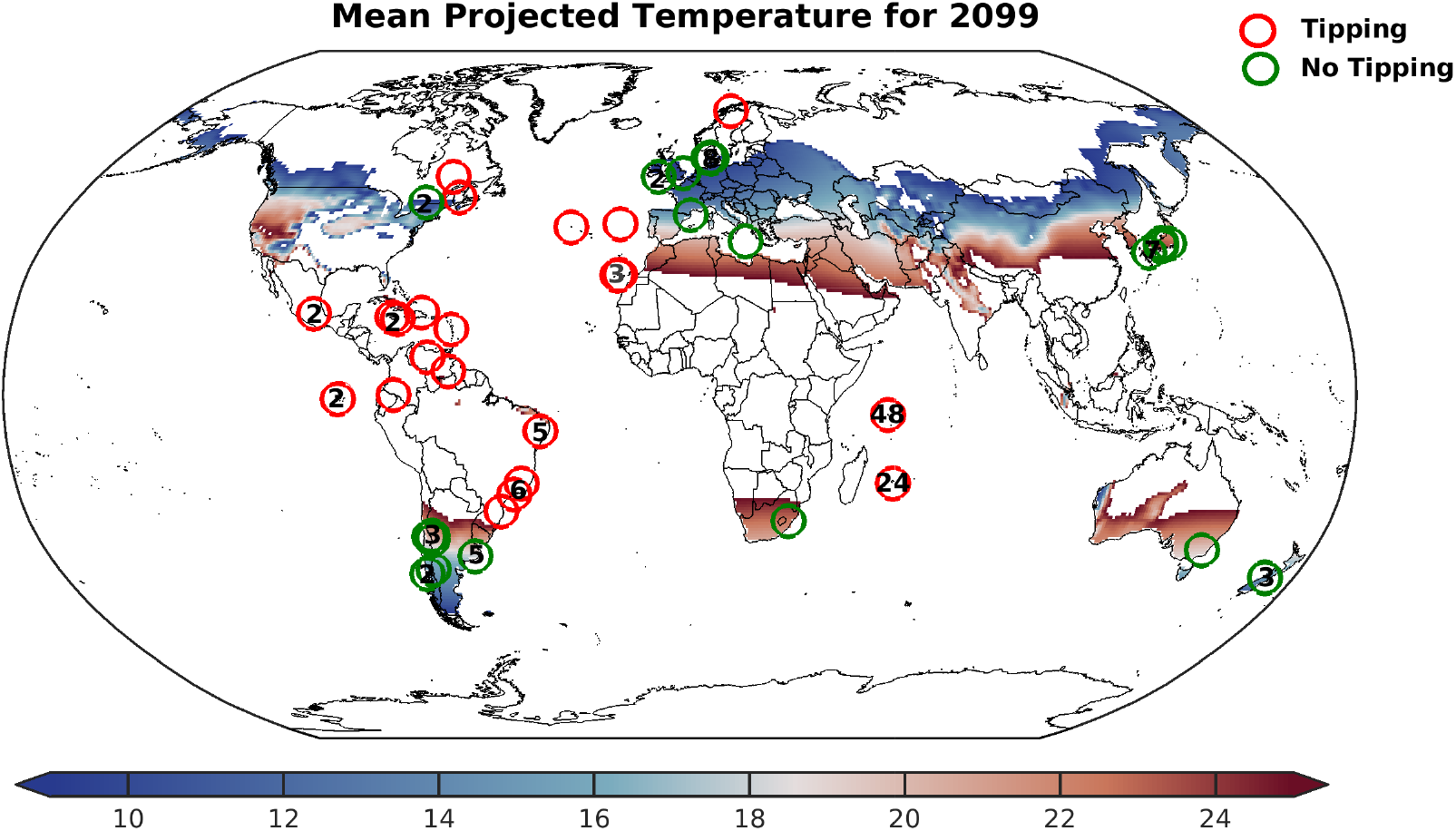
Chances of tippings and No tippings in networks based on location-wise future projected global temperature: The figure shows 139 plant-pollinator networks plotted at their respective locations. Color codes are representative of the regions experiencing temperatures in the optimum range (9 – 25°*C*) in the color-bar in 2099. Insets (green) denote the 37 networks that do not undergo tippings being present in regions experiencing optimum temperature and 102 networks (red) that can undergo tippings in 2099 on following socio-economic pathway (SSP370, CMIP6).

We characterize our results in Fig. 7 as a rudimentary tool providing indications to study the networks exhibiting risk of tipping in greater detail. It may be noted that, networks which undergo tipping in the present simplified set up, the study of dynamics of such networks should be of the highest priority for further study. One should not however conclude that such networks would definitely undergo a critical transition in future.

Nevertheless, envisioning the increased rate of global warming and predicting the fate of the real mutualistic networks, formulation of practical management policies is of a dire need to avoid frightful consequences. The development of realistic mitigation policies requires model parameters varied in a range that concede with the biological constraints. Taking care of such constraints, only a few management strategies are realizable in effect. Some of the cost-effective, viable principles involve maintaining the abundance of an influential pollinator or abate the decay rate of the same to a bare minimum (Jiang et al 2019). Here, we demonstrate how inherent network structures interplay with the degree of warming to avoid tippings.

We study how maintaining the abundance of the most generalist pollinator to a constant value can aid in the early recovery of the mutualistic community, which would have otherwise remained extinct. We have shown our result taking four different networks; network A (*M_PL*_061_33, *S_A_* = 6, *S_P_* = 2, *NODF* = 0), network B (*M_PL*_061_14, *S_A_* = 11, *S_P_* = 6, *NODF* = 0.25), network C (*M_PL*_006, *S_A_* = 61, *S_P_* = 17, *NODF* = 0.52), network D (*M_PL*_059, *S_A_* = 13, *S_P_* = 13, *NODF* = 0.84). Simulations of 4 different networks reveal that nestedness has a critical role in steering the recovery of species at higher temperatures. We observe that without any mitigation, the networks do not recover beyond 32°*C* irrespective of their underlying structure. On fixing the abundance of the influential pollinator at a abundance of 0.2, these systems with distinct underlying structures recover for the above networks. The point of recovery is a latent function of the complex interaction of the habitat temperature and structural properties. As the mean habitat temperature is increased beyond 32^0^*C*, we find that networks with high nestedness recover at a decreased *γ*_0_ value. At 40^0^*C*, the network “A” with nestedness (NODF) value 0 does not recover despite preserving an abundance of the generalists’ species while networks “B” and “C” can be retrieved on further increasing *γ*_0_ values respectively, and network “D” with value 0.84 recover at *γ*_0_ = 1.9 (Fig. 8). In Figs. 8(e)-8(h), we observe that at 40°*C*, when there is a surge in the species decay rate, the threshold abundance of the generalist species requires to be upraised. An increase in the fixed abundance of the generalist will aid in recovering the community at a reduced *γ*_0_ value. This recovery point is further enhanced for a highly nested network, yet relationships are highly nonlinear. An important observation includes the fact that an average *γ*_0_ for low to moderately nested networks is essentially greater than or comparable to intraspecific competition (*β_ii_* = 1) promotes recovery.

**Figure 8.**
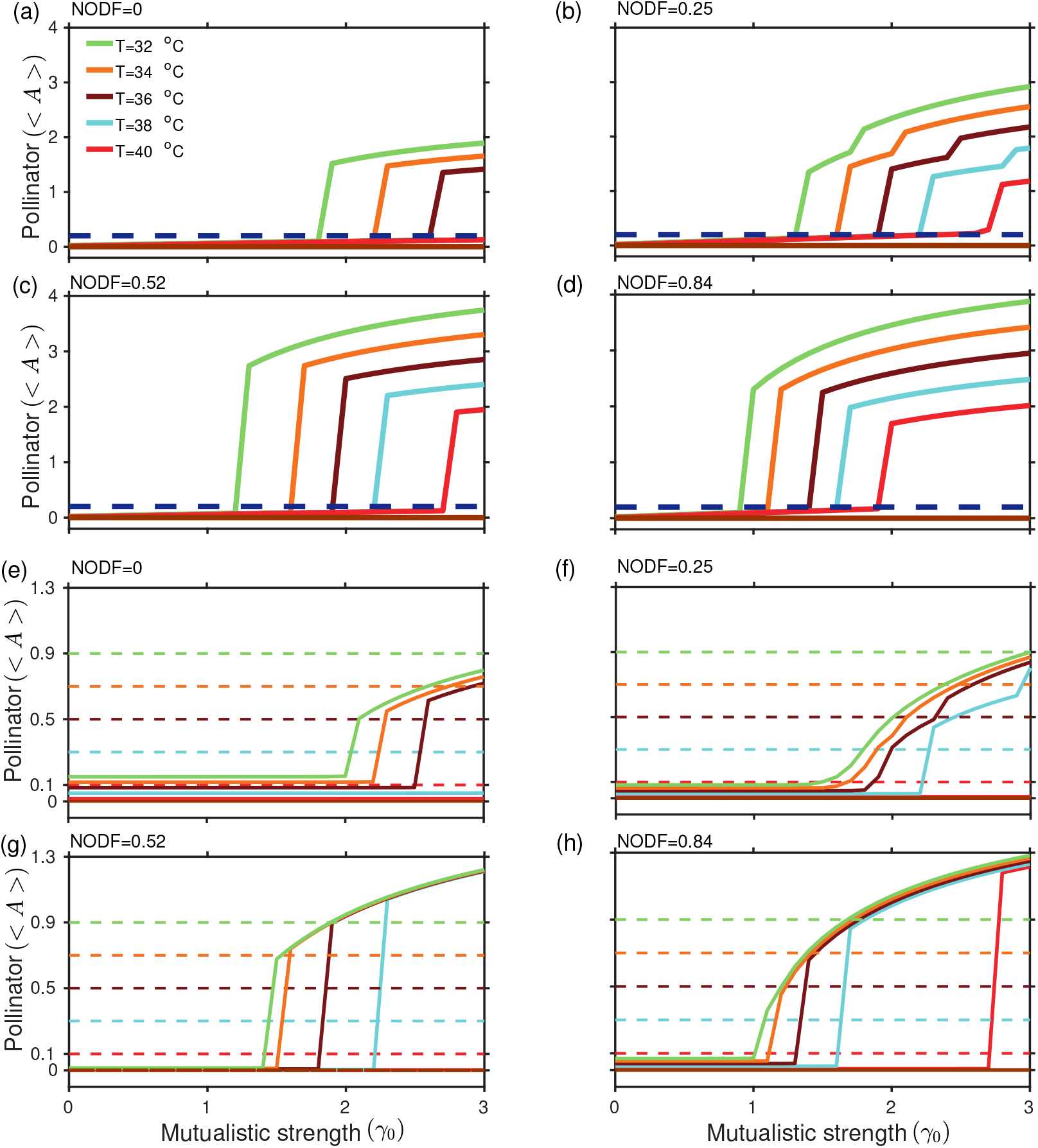
Role of network structure in managing tippings in mutualistic networks: (a)-(d) Abundance management in 4 different networks in the increasing order of nestedness (NODF). The dashed line (blue) denotes the fixed abundance of the generalist species as a management strategy, and the solid line (maroon) denotes the average abundance of species in the absence of management strategy at and beyond 32°*C*. The average abundance of the community is plotted at various temperatures above 32^0^*C*. Systems with higher nestedness experience early recovery for all the temperatures in the range (32 – 40)°*C*. At 40°*C*, the network A (NODF=0) fails to recover with the generalist species abundance fixed at 0.2, whilst network B (NODF=0.25), C (NODF=0.52), and D (NODF=0.84) recover. The network D undergoes the fastest recovery. (e)-(h) Dashed line denote the abundance of the generalist species fixed at abundances 0.1, 0.3, 0.5, 0.7, and 0.9, respectively, and the recovery of the community at 40° C are correspondingly plotted in the same color with a dashed line. The solid line (maroon) represents the average abundance of the realized networks without any control. At 40°*C*, the network A (NODF=0) fails to recover with the generalist species abundance fixed at 0.3 and less, whilst network B (NODF=0.25) and C (NODF=0.52) do not recover with the generalist species abundance fixed at 0.1 and less, but network D (NODF=0.84) recovers with the generalist species abundance fixed at any value greater than or equal to 0.1. We observe fixed threshold abundances of the generalist pollinator at 40°*C* for networks A, B, C, and D, which aid in the system’s recovery.

Another acceptable strategy is where the pollinator with the highest degree (i.e., generalist pollinator) is targeted, and factors leading to its decay are minimized by setting its decay rate (*k_i_*) to 0.01, while other pollinators have high decay rate. Figure 9 represents the average community trajectories at different temperatures above 32°*C*, at which the networks experience global extinction. Nestedness plays an equivalent vital role in expediting recovery at high temperatures with profound effect at the extreme. Despite the intraspecific competition, as the level of nestedness increases, pollinators interact more with generalist plants. Pollinators forming a part of a nested network tend to hold back the community on the verge of collapse and promote early recovery. Therefore viable control principles should also involve adaptive strategies to increase nestedness of mutualistic communities alongside reducing environmental stress.

**Figure 9.**
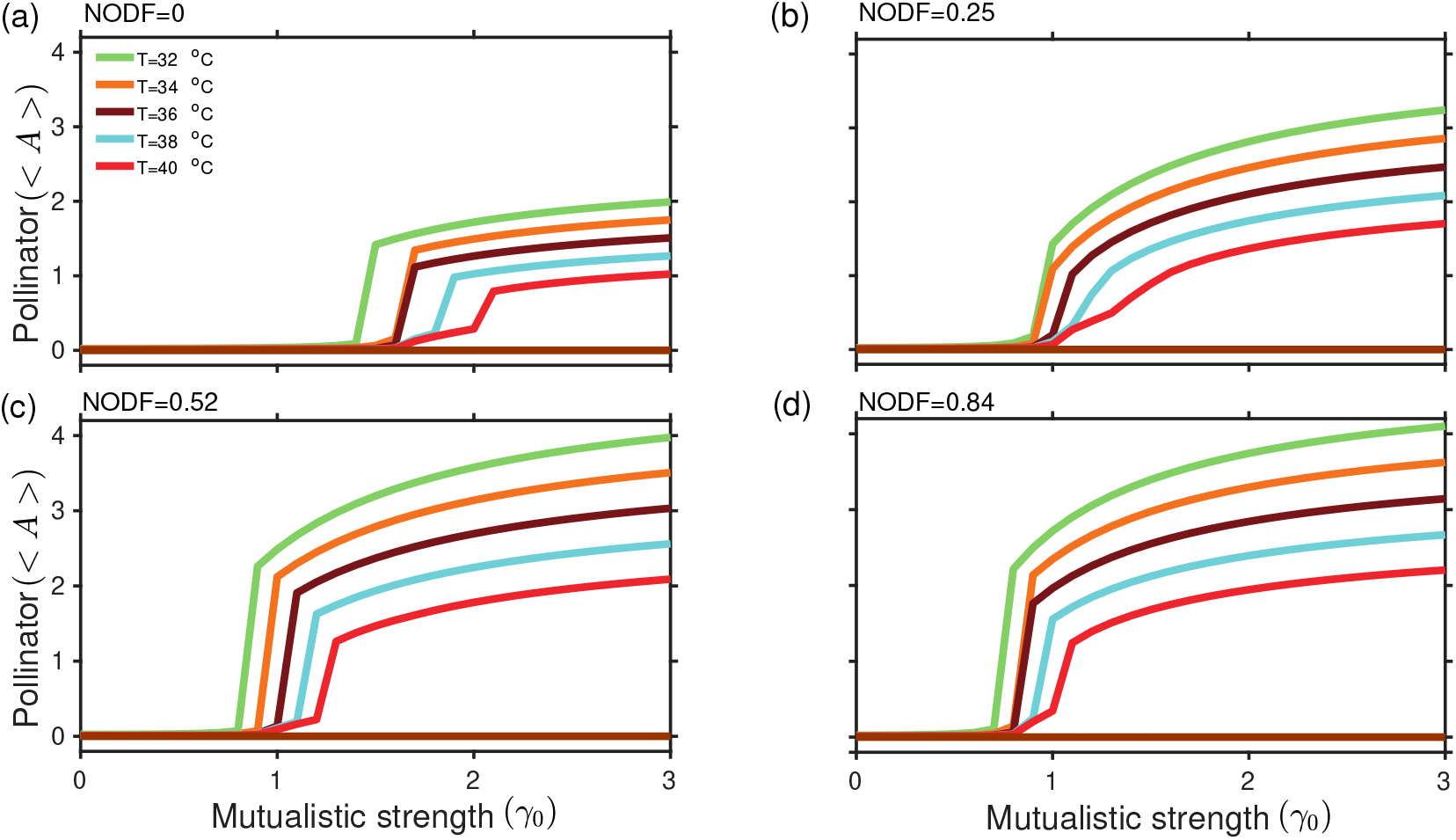
Managing tipping by minimizing decay rate of the generalist species: At various temperatures above 32°*C*, the death rate of the generalist species for networks A, B, C, D is set at 0.01. Solid lines (maroon) represent species abundance in the absence of management strategy. For network A (NODF=0), recovery is considerably delayed at the extreme temperature (40°*C*).

## Discussion

Studies concerning the effect of species’ individual thermal responses of life history traits on ecological networks, such as mutualistic networks, remain less stated. Collected data of plants and their pollinating insects from Illinois, USA, in the late 18th century and revisited back in 2010 reveal loss in pollinator functioning and reduced adherence to their mutualistic counterparts (Burkle et al 2013). Such information impel the study of network dynamics as species’ individual growth rate, birth rate, and handling time are known to alter with the degree of warming. Owing to the fact that recovering species/community post collapse demands lowering temperature upto a few degrees or increasing mutualistic strength among mutualists, our explorations revolve around the variations in latter with network structural properties.

While an increase in temperature can push mutualistic communities at the brink of collapse (Fig. 1), increasing *γ*_0_ aids in species rescue and prevent community collapse (Figs. 1(d)-(f)). We find an interesting yet alarming fact, as *γ*_0_ is reduced, an increase in temperature beyond 28°*C* sets off catastrophic transitions (Fig. 2). A hysteresis loop is built up, which explains the increased net effect of mutualistic species on one another at high temperatures. These results are generalized and hold good for networks of varied size and structural properties (*SI Appendix*, Section S1, Figs. S1.4,S1.5). In context to the robustness of our results although, for simplicity we assume equal coefficient and symmetric functional response curves - qualitative results remain similar on considering functional response curves distributed randomly around a mean with a given variance (*SI Appendix*, Section S1, Fig. S1.2) or skewed species response rates. While perturbations in species functional rates produce fluctuations in the abundance of pollinators without affecting the tipping point, skewed response curves shift the tipping temperature for a considered *γ*_0_ correlated with the shift in the optimal temperature. Likewise, incorporating non-zero interspecific competition (*β_ij_* ≠ 0), the requisite *γ*_0_ that aids in preventing species extinction increases (*SI Appendix*, Section S1, Fig. S1.3). If one assumes *α_i_*(*T*)/*β_ii_*=*K_i_*(*T*), where *K_i_*(*T*) is the temperature dependent carrying capacity, then it opens the possibility of considering a variety of shapes for *α_i_*(*T*) following the study of (Uszko et al 2017). Interestingly, mutual dependence is a two-sided coin. At high habitat temperatures, as *γ*_0_ decreases, a sudden decline in plant abundance can trigger the extinction of pollinators and vice versa. However, an increase in mutualistic strength between the interacting species can aid in the recovery of both. Although, we envision tipping, gaining insights into the dynamics of the higher dimensional network model are rather difficult. Under this backdrop, study of three minimal network models provide useful inferences–plunging more plants into the network with fixed number of pollinators can be a management policy to prevent pollinator collapse at the face of climate warming (*SI Appendix*, Section S1, Fig. S1.6).

In a deteriorating environment, as the driver of pollinator decline increases and node or link loss becomes inevitable, preserving the key species can prevent community collapse. Removal of a fraction of generalists (Fig. 4) have a more harmful impact on the network compared to their specialist counterparts or random removals of plant or pollinator (*SI Appendix*, Section S1, Fig. S1.11-S1.13) at high temperatures, since it is connected with more number of species. Designing conservation strategies to prevent network collapse demands lowering the degree of climate warming or maintaining the requisite network structural properties. Network collapse may be averted or delayed by preserving generalist species.

Our results have implications in fostering network resilience at high temperatures for the 139 real plant-pollinator networks. An intriguing result is that high nestedness (and low modularity) delays collapse or allows the system to recover early after being perturbed at extreme temperatures (37° – 40°*C*) (Fig. 3). This allows us to claim that maintaining the optimal structure can confront issues related to network collapse due to anthropogenic stress. Managing networks at a high temperature by maintaining constant abundance or minimizing the decay rate of the most generalist pollinator is more straightforward for networks with higher or moderate nestedness values as they recover even for a low mutualistic strength. With regard to the feasibility of such techniques to empirical networks, artificially experiments need to be performed in the laboratories. One such experiment has successfully been able to increase the connectance of plant-pollinator networks. The field experiment increased the connectance by improving attractiveness of plant species in a bee-plant network (Russo and Shea 2017). In the absence of any management policies, networks irrespective of their structural properties do not recover at 40°*C* (Fig. 8, and Fig. 9). We also demonstrate this by the stability analysis of the reduced temperature-dependent two-dimensional model (*SI Appendix*, Section S2, Fig.S2.2). As observed, the region of stability reduces i.e., the system remains unstable for a large range of *γ*_0_. Hence small environmental perturbations can push the system to an alternate extinction state. It may be noted that the interrelations between nestedness and modularity can differ across different mutualistic networks, and modularity does not always have a negative effect unlike our set of empirical networks (Fortuna et al 2010).

Our study is limited by the fact that the mutualistic strength *γ*_0_ is not a function of temperature. Primarily, it is due to lack of empirical evidence. Nevertheless, *γ*_0_ indirectly accounts the effect of changing temperature via the abundances of plants and pollinators. Therefore, our study provides with a framework on which to build a more realistic model into the future and has the potential of answering questions related to conditions that support partnership and improved predictions at the face of global warming. While our study revolve around the effects of gradual temperature change on mutualistic plant pollinator communities; there remains a few open questions. An interesting future direction includes studying the effects of extreme climate events such as floods, droughts, and erratic heat wave days over a period of time on an assembly of mutualistic community. This may be investigated by incorporating infrequent sudden large stochastic perturbations in the mean habitat temperature of the system (Ummenhofer and Meehl 2017, Thibault and Brown 2008). Another promising direction is to study the temperature driven network dynamics and the evolution of species while they exhibit niche based interactions with their mutualist counterparts (Cai et al 2020). The modified framework would also allow understanding the effects due to various factors such as anthropogenic changes and invasive alien species. As studied by (Valdovinos 2019), acknowledging adaptive behavior can alter stability relations. Considering adaptive foraging of pollinators with increase in climate warming by modelling the adaptation of animal species foraging preference on plant via a differential equation within the present framework will be of substantial ecological importance. One may consider different temperature response curves for juvenile and adult species and study under the framework of stage-structured mutualism (Nakazawa 2020). This is important yet challenging and requires visiting sites, collecting data, and finding the functional curves, which might demand continuous monitoring of sites at regular intervals. Our study tells that behaviors at the individual level beget collective system-level consequences. Once network sites are revisited, and different species functionality with temperature variations are closely monitored, our results can be tested further to provide valuable insights to other real ecological networks.

## Acknowledgments

S.D. acknowledges the Ministry of Education (MoE), Govt. of India for Prime Minister’s Research Fellowship (PMRF). P.S.D. acknowledges financial support from the Science & Engineering Research Board (SERB), Govt. of India [Grant No.: CRG/2019/002402]. The authors thank Christopher F. Clements for helpful discussion on the model, and Sourangsu Chowdhury for helpful discussion on the CMIP6 data. Output from the CMIP6 models from https://esgf-node.llnl.gov/projects/cmip6/ is greatly acknowledged.

