## SUPPORTING INFORMATION for "Rising Temperature Drives Tipping Points in Mutualistic Networks^†^"

#### S1: Temperature driven dynamics of various real plant-pollinator networks

The species thermal response curves considered for studying the dynamics of plant-pollinator networks are shown in Fig. S1.1(a)-S1.1(c). We show that our results are robust to the effect of perturbations in species functional responses (Fig. S1.2) and non-zero interspecific competition (Fig. S1.3). We present results for networks of varied dimension and structural properties (see Table. S1.1). Our results are generalizable and hold good for a wide array of mutualistic networks (Fig. S1.4, Fig. S1.5) . Irrespective of the networks size and structure, hysteresis is vivid across all the considered 16 networks (Fig. S1.5). Fig. S1.6 presents the dynamics of 3 minimal models for varying temperature at different mutualistic strengths ($\gamma_0$). We also present the role of other structural properties viz connectance and network size on tipping point in mutualistic networks (Figs. S1.7 ,S1.8). The effect of trade-off on temperature is also consistent across the majority of networks with a more profound impact of high $\delta$ values on tipping at high temperatures (see Fig. S1.9). We also report the correlation between nestedness of the the 141 plant pollinator networks and their point of collapse ($\gamma_0$), at different temperatures (see Fig. S1.10). We study the effects of removal of fraction of specialists (Fig. S1.12) and random removal of fraction of plant and pollinators for network $M\_PL\_006$ on community collapse in comparison to removal of their generalist counterpart (see Fig. S1.11, S1.13). Table. S1.2 presents details of species in network $M\_PL\_006$ and a count of their presence in other networks used for the above study.

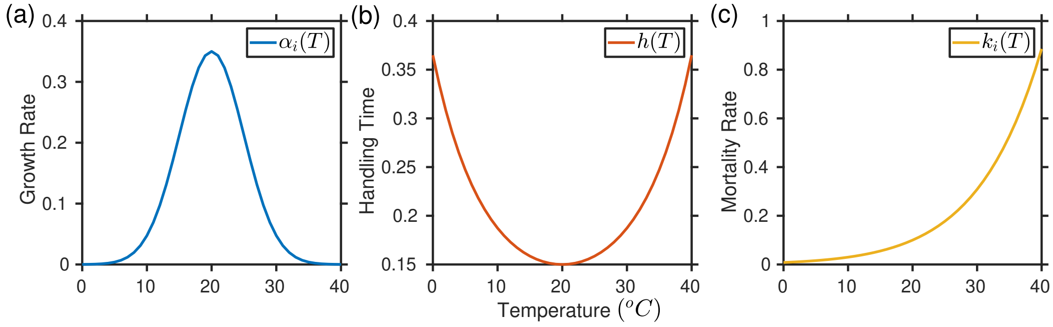


**Figure S1.1. Thermal response curves of species biological traits**: Effects of rising temperature on the birth rate (blue), handling time (red), and death rate (yellow) of species. The parameter values are $\alpha_{opt} = 0.35$, $T_o = 293K(20^oC)$, $\sigma_\alpha = 5$, $h_{opt} = 0.15$, $\sigma_h = 15$, $k_{opt} = 0.1$, and $A_K = 10^4$.

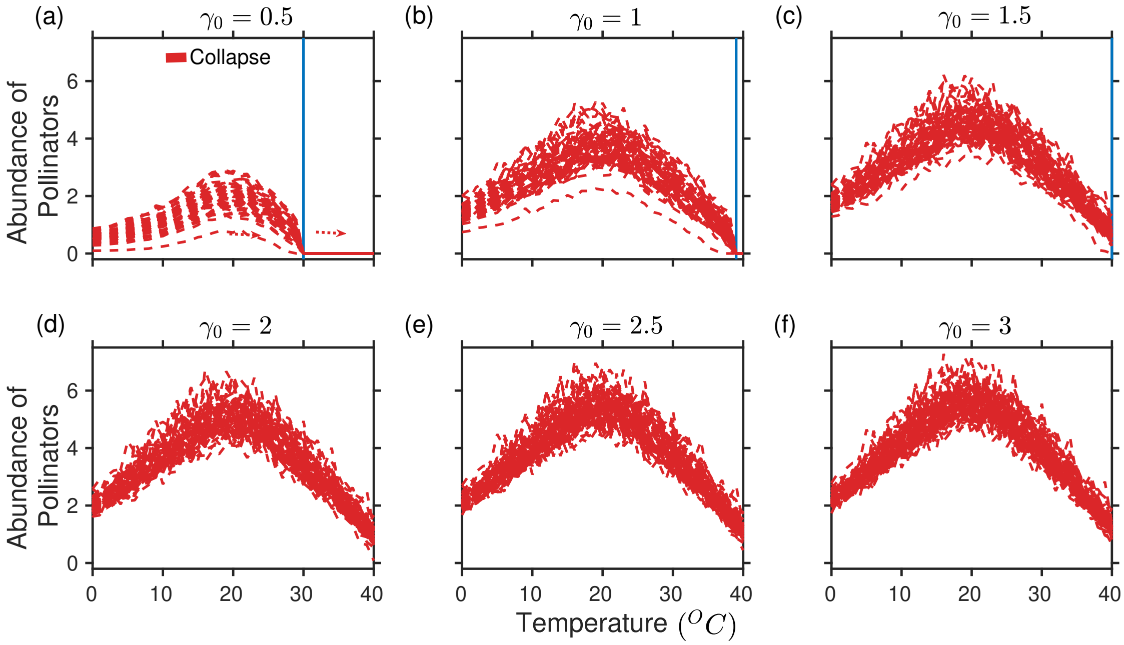


**Figure S1.2. Effect of heterogeneity in the values of $\alpha(T)$, $h(T)$ and $k(T)$ on driving tipping points**: (a)-(b) On increasing temperature in the range $(0 - 40^oC)$, at $\gamma_0 = 0.5$ to $\gamma_0 = 1.0$, the system encounters a catastrophic transition. (c) At $\gamma_0 = 1.5$ the system undergoes gradual decline in density. (d)-(f) At and beyond $\gamma_0 = 1.5$, community collapse is averted, despite there is a gradual drop in abundance. Heterogeneity in the temperature dependent parameters lead to fluctuations in abundances although produce no additional effect on the system dynamics. The blue vertical line marks the occurrence of a critical transition. The above result is obtained for the network $M\_PL\_006$ with $S_A = 61$ and $S_P = 17$ (for details see "http://www.web-of-life.es/"). The parameter values are $\beta_{ii}^A = \beta_{ii}^P = 1$, $\delta = 0.5$, $\mu_A = \mu_P = 10^{-4}$, and the other parameters $\alpha_i^A$, $\alpha_i^P$, $k_i$, $h$ are distributed about a mean same as the rate value of respective response function and a variance of 10% about the mean.

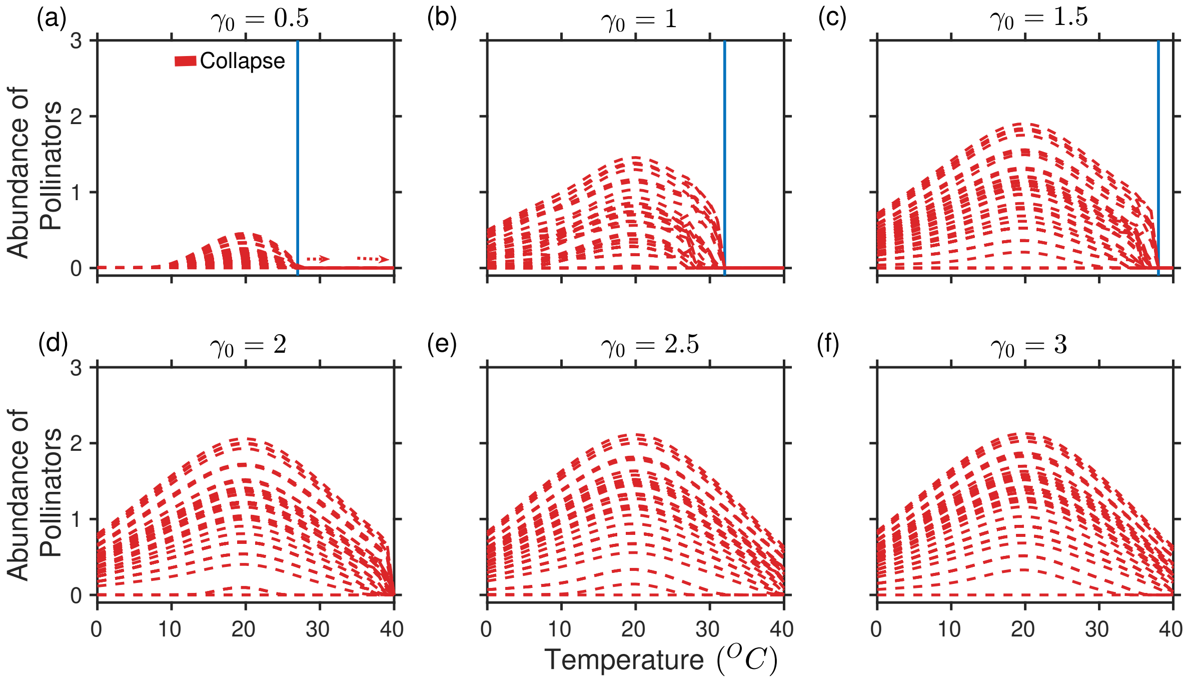


**Figure S1.3. Temperature driven network dynamics considering nonzero interspecific competition**: (a)-(e) On increasing the temperature in the range $(0 - 40^o C)$, for $\gamma_0 = 0.5$ to $\gamma_0 = 2$, the pollinator populations encounter catastrophic transitions. (f) At or beyond $\gamma_0 = 2$, sudden community collapse is averted, despite there is a gradual drop in the abundance of pollinators with increasing temperature. Incorporating non-zero interspecific competition causes reduced pollinator abundance demands comparatively higher $\gamma_0$ values to evade species extinction. The blue vertical line represents the occurrence of a critical transition. The above result is obtained for network $M\_PL\_006$ with $S_A = 61$ and $S_P = 17$ (for details, "http://www.web-of-life.es/"). The parameter values are $\beta_{ij}^A (i \neq j) = \beta_{ij}^P (i \neq j) = 0.05$, $\beta_{ii}^A = \beta_{ii}^P = 1$, $\delta = 0.5$, $\mu_A = \mu_P = 10^{-4}$, and the other parameters $\alpha_i^A$, $\alpha_i^P$, $k_i$, $h$ are obtained from their respective response function and are considered equal for simplicity.

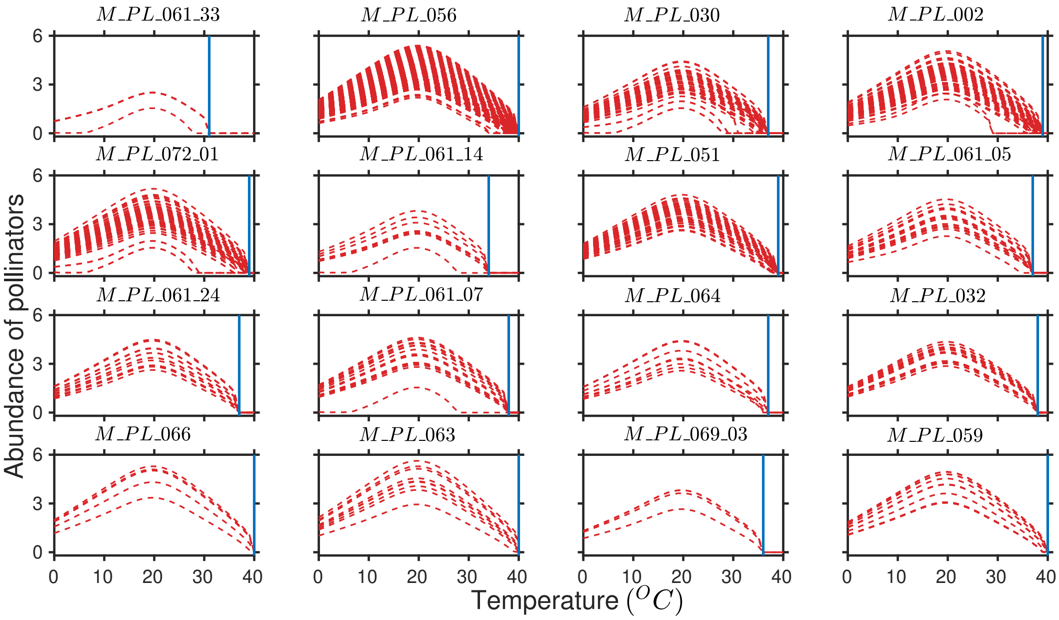


**Figure S1.4. Higher temperature can trigger catastrophic transition across mutualistic networks**: On increasing temperature in the range ($0 - 40^0C$), at $\gamma_0 = 1$ the system undergoes a catastrophic collapse. The blue vertical line represents the occurrence of a critical transition. We have shown this result considering 16 networks with different nestedness values chosen from the database www.web-of-life.es (see Table. S1.1 for further details). Results are consistent across all the networks.

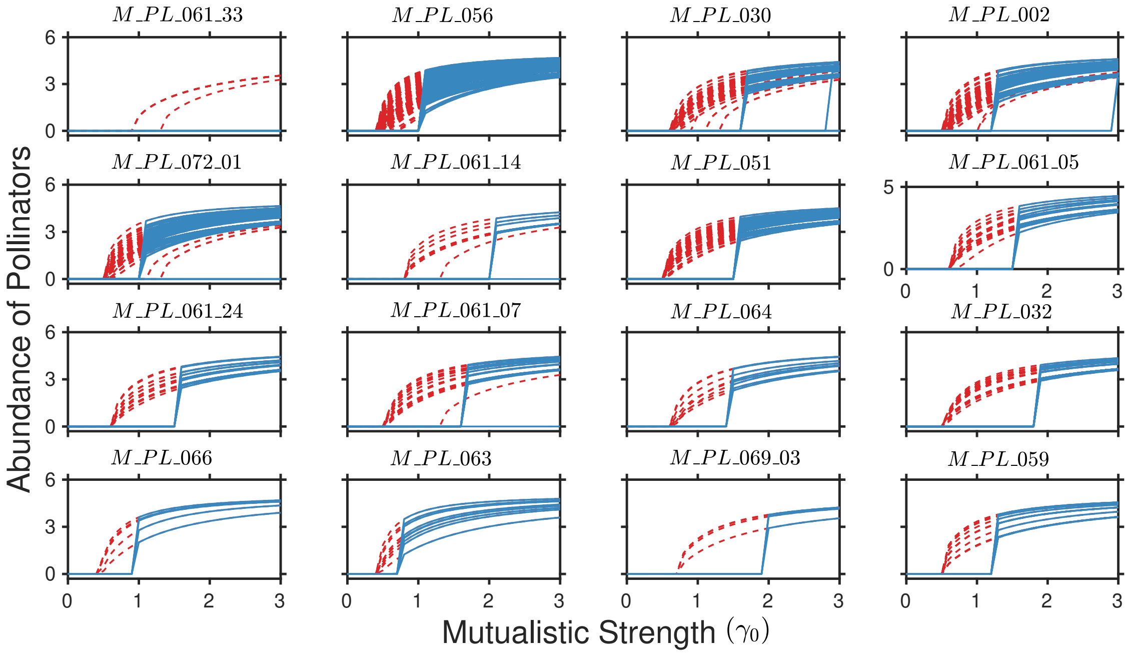


**Figure S1.5. Catastrophic collapse in mutualistic networks for variation in mutualistic strength ($\gamma_0$**: On varying the mutualistic strength $\gamma_0$ in the backward direction $(3 - 0)$, at high temperature ($30^oC$), the system encounters a catastrophic transition. In the forward direction, the system can be recovered by reducing environmental stress (varying $\gamma_0$ from $(0-3)$). Point of collapse and recovery are different, indicating the formation of a hysteresis loop observed for all networks at extreme temperatures. We have shown this result considering 16 networks with different nestedness values chosen from the database www.web-of-life.es (see Table. S1.1 for further details). Results are consistent across all the networks.

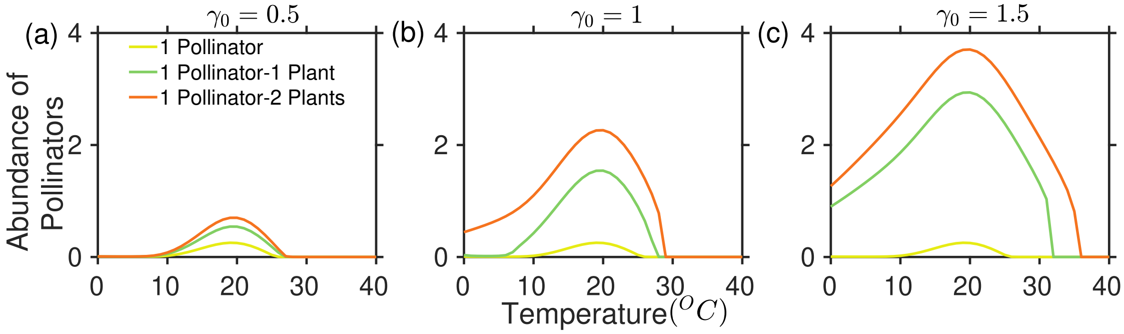


**Figure S1.6. Effect of increasing temperature on the dynamics of minimal mutualistic network models**: On plotting the pollinator abundance against temperature gradients for (a) $\gamma_0 = 0.5$, (b) $\gamma_0 = 1$, and (c) $\gamma_0 = 1.5$, we observe that adding plants into the mutualistic network aids in delaying collapse of the pollinator while also increasing its abundance. Effects are more profound for higher $\gamma_0$ values. In each sub-figure, color of a trajectory denotes the pollinator abundance in one of the three minimal models. All the other parameter values are same as Fig. 1 (main paper).

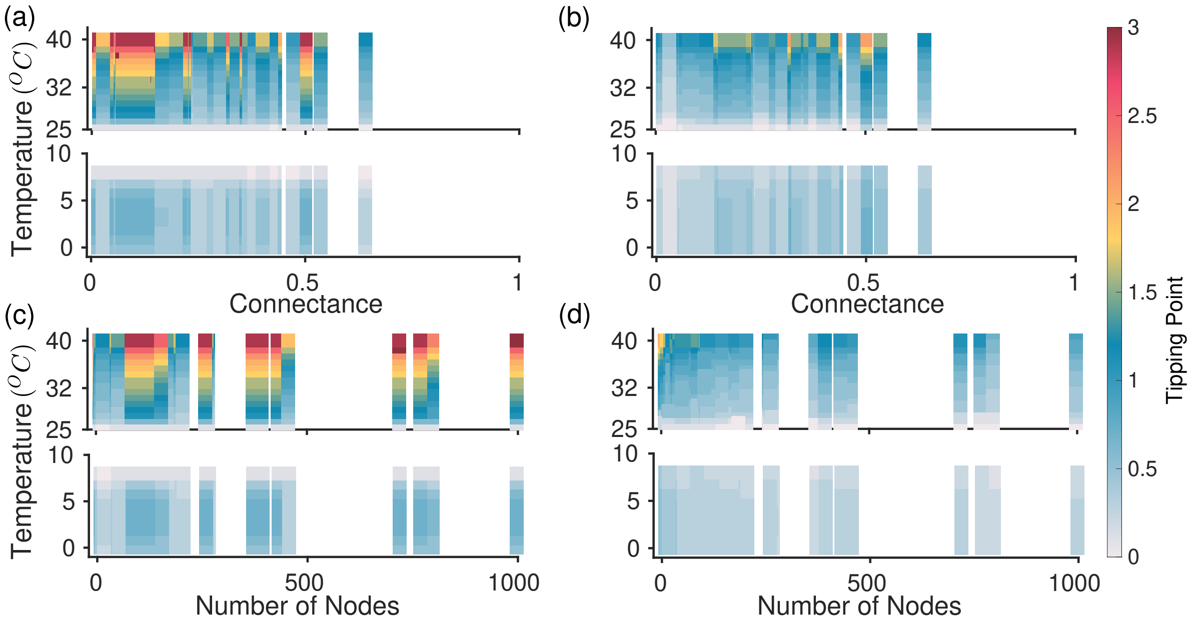


**Figure S1.7. Role of network properties (i.e., connectance and number of nodes) under varied degree of warming in affecting a tipping point**: (a) First point collapse of pollinator populations for 141 real plant-pollinator networks plotted across their respective connectance values. At higher temperatures, more connected networks undergo collapse at a considerably lower $\gamma_0$ value (which marks the tipping point and presented in the color bar). (b) Final point collapse of the plant-pollinator community plotted across their respective connectance values. (c) First point collapse, and (d) Final point collapse of all the networks are plotted across their respective number of nodes. The colors in the color bar correspond to the $\gamma_0$ values in the range $(0 - 3)$, at which the system undergoes a first (a,c) and final point collapse (b,d). The results are averaged over 100 independent simulations.

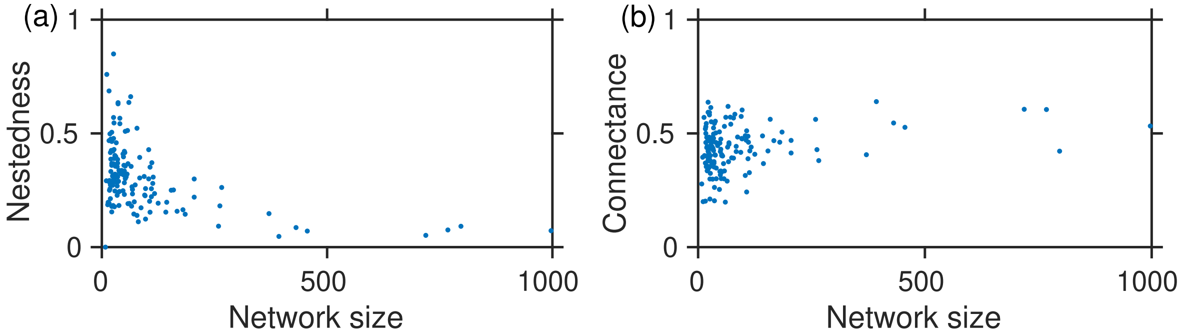


**Figure S1.8. Correlation between network properties and network size**: For the 141 real plant-pollinator networks both (a) Nestedness and network size (b) Connectance and network size show no prominent inter-relation trends.

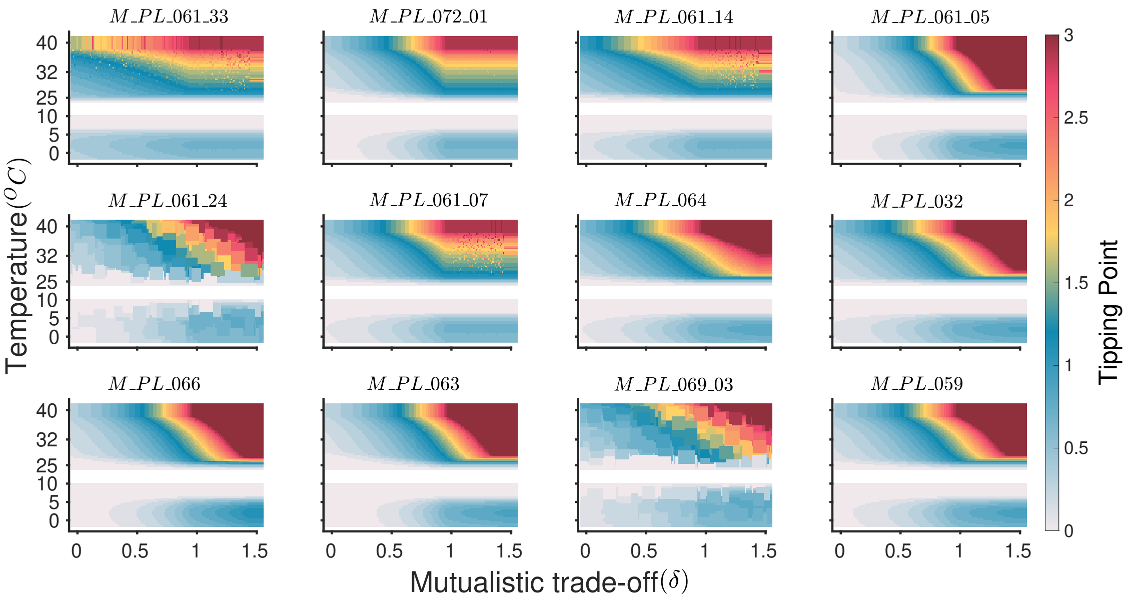


**Figure S1.9. Effect of trade-off on the temperature driven network dynamics**: For each of the 12 observed mutualistic networks, this figure illustrates that the point of collapse is attained much earlier for high mutualistic trade-offs at high temperatures. For optimum temperature, collapse can be avoided for any mutualistic trade-off. With an increase in the trade-off ($\delta$), the interdependence among species grows at high temperatures resulting an early (high $\gamma_0$ values) community collapse. As the temperature rises, the vulnerable species that are the first to get extinct also incite sudden transitions in the other species, rapidly approaching the collapse point. At higher temperature effects are significantly more profound specifically for $\delta$ beyond 1. We did not consider the intermediate temperature range $9 - 25^oC$ as the system does not undergo collapse in this range.

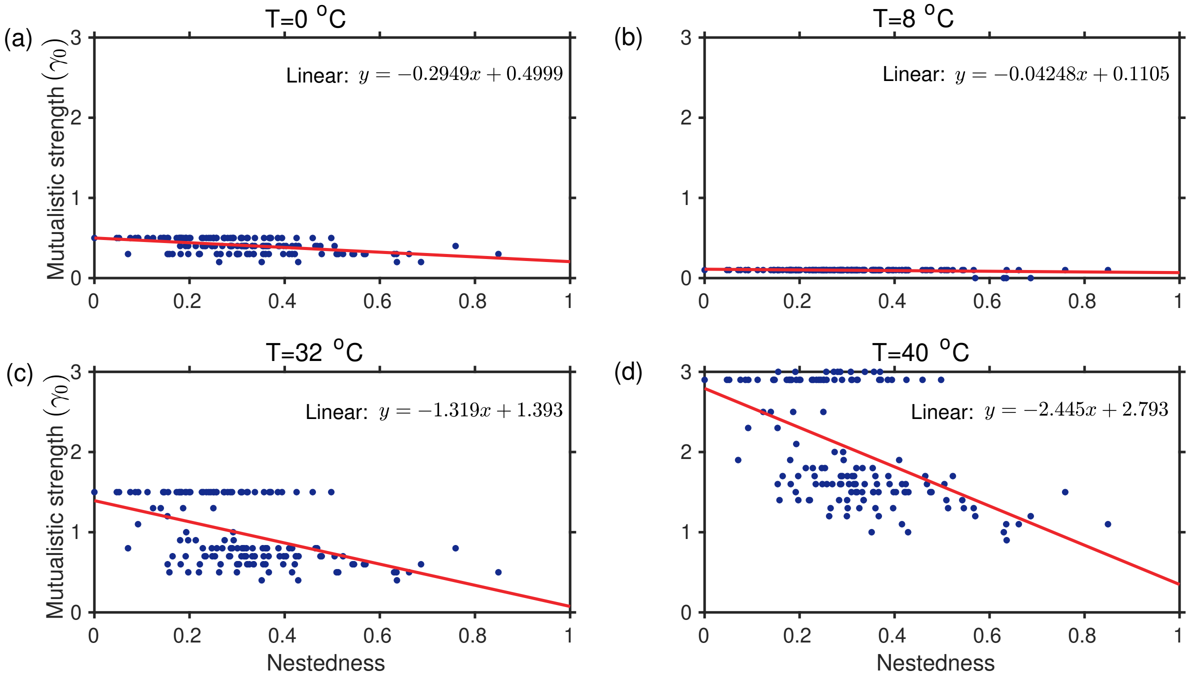


**Figure S1.10. Relation between nestedness and tipping points**: For each of the 141 real mutualistic networks, the results depict that for a network with low NODF value, systems move more rapidly towards tipping than a network with high NODF values. This can be interpreted as tipping points and nestedness are negatively correlated at high temperatures beyond the optimum. The negative correlation is much stronger at high temperatures. (a)-(d) Slope of the fitted lines are $-0.29$, $-0.04248$, $-1.319$, and $-2.445$, respectively.

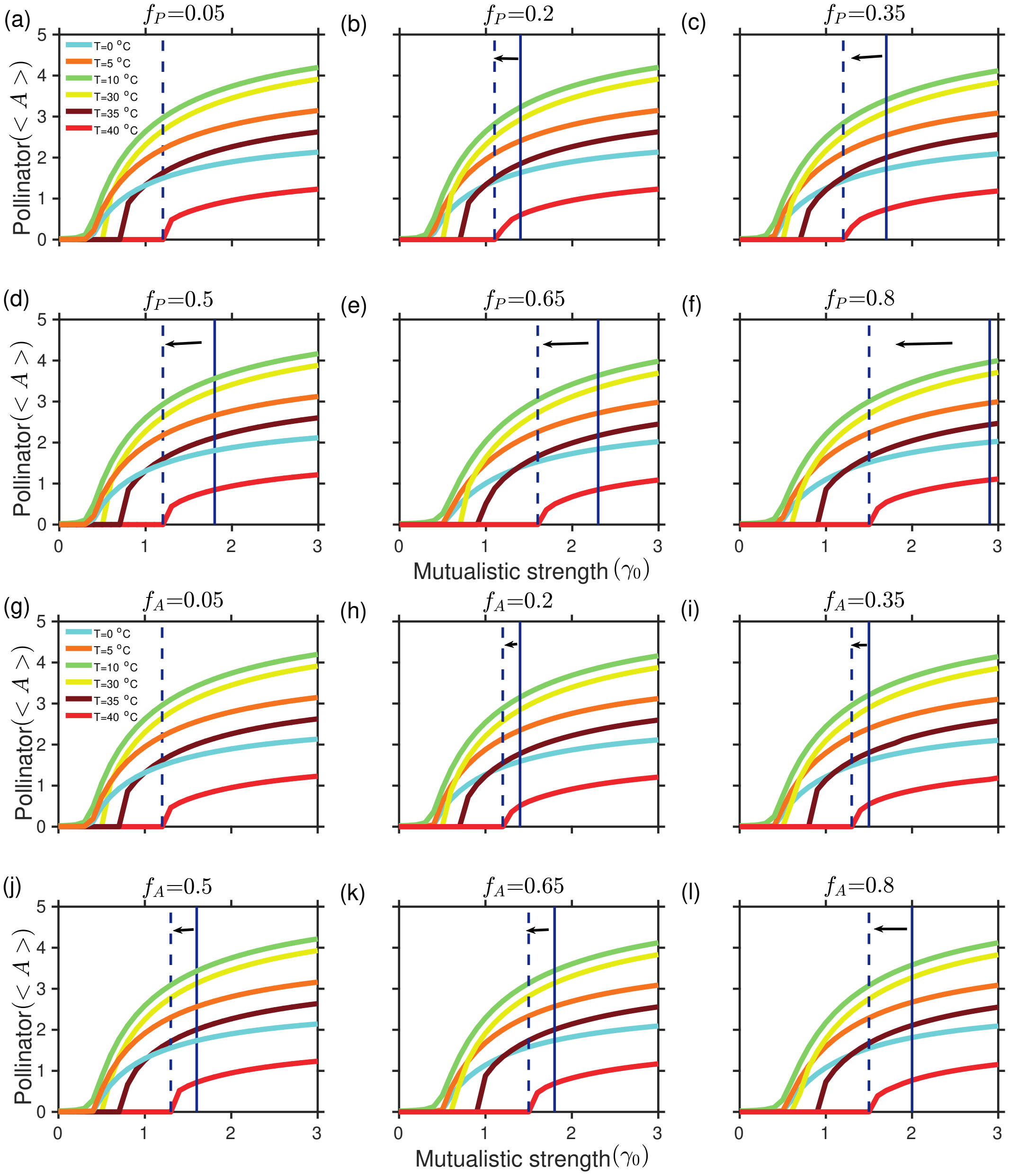


**Figure S1.11. Effects of random removal of plant and pollinator for varied temperature regimes**: Panels (a)-(f) and (g)-(l) depict the decline in the average abundance of the pollinator community as a fraction of plant and pollinator are removed from randomly chosen nodes. As the mutualistic strength $\gamma_0$ is gradually decreased from 3-0, the point of collapse precedes as $f_P$ and $f_A$ are increased further. As observed in (f) and (l), on the removal of random plant or pollinator species remarkable difference is observed in tipping at high temperature ($40^0C$) for an increase in $f_P$ and $f_A = 0.8$. Highly connecting species removal has a more prominent effect on the collapse of the pollinator community than random removal of species as indicated by the distance between solid and dashed vertical lines for respective $f_P$ and $f_A$ values. The solid line in all the panels (a)-(l) represents the point of collapse on the removal of a fraction of plants ($f_P$) and pollinators ($f_A$) in decreasing order of their degrees (see Fig5, main text) and the dashed line represents the collapse when the corresponding fraction of plant and pollinators are removed randomly.

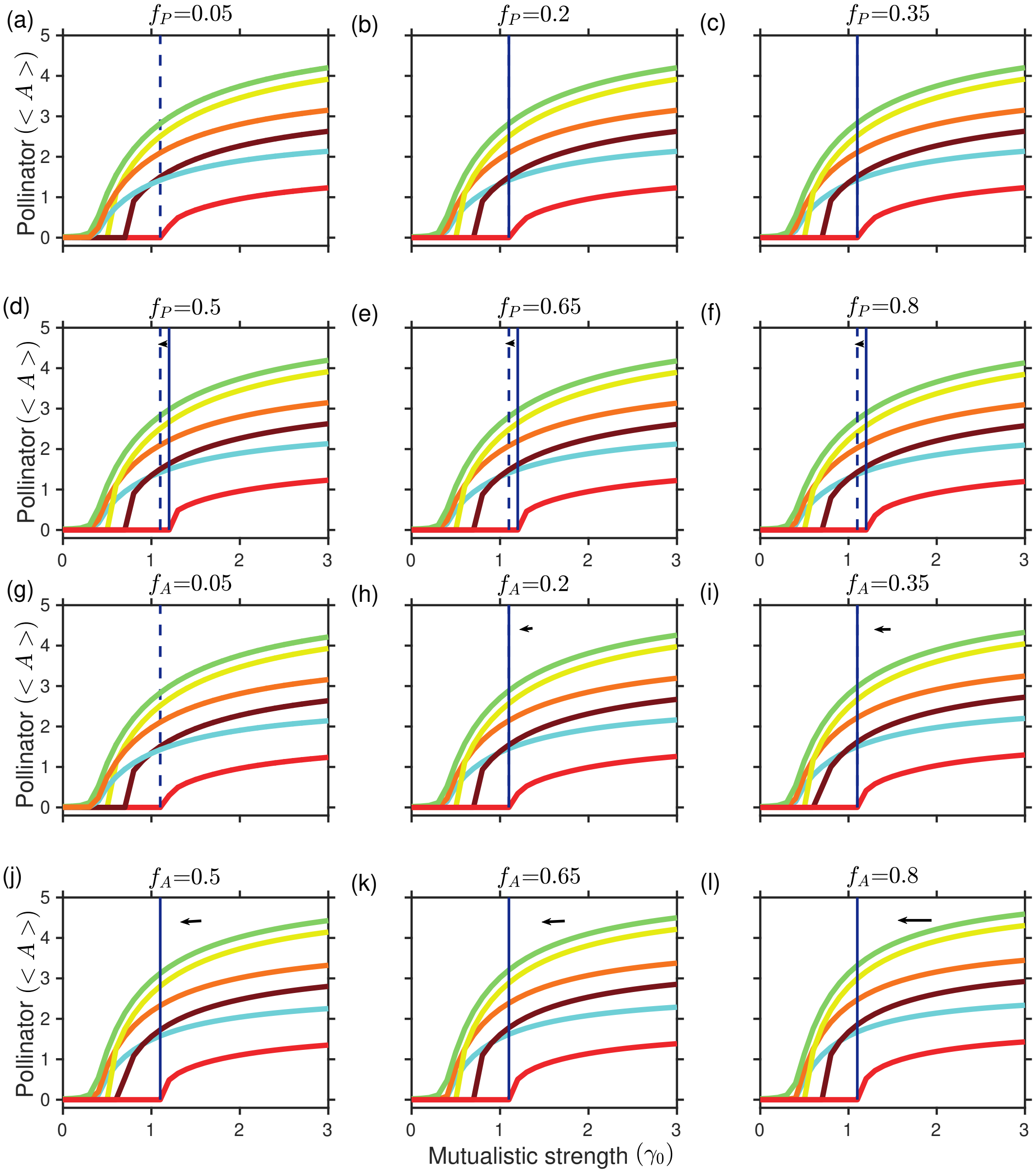


**Figure S1.12. Effects of specialist plant and pollinator loss for varied temperature regimes**: Panels (a)-(f) and (g)-(l) depict the decline in the average abundance of the pollinator community as a fraction of specialist plant and pollinator are removed in increasing order of their degree at temperatures ranging over the interval $0 - 40^oC$ (denoted by different colors in the legend). Gradually decreasing the mutualistic strength $\gamma_0$ from 3-0, the point of collapse precedes as $f_P$ and $f_A$ are increased further. As observed in (g)-(l), on removal of specialist pollinator species no prominent difference is observed in tipping at high temperature ($40^0C$) for an increase in $f_P$ and $f_A = 0.8$. Highly connecting species removal has a more prominent effect on the collapse of the pollinator community than random and less connected removal of species. The dashed lines in (a)-(l) represent the point of collapse for $f_P = f_A = 0.05$. The solid lines in (b)-(f) and (h)-(l) represent the point of collapse for different values of $f_P$ and $f_A$ as mentioned on top of each sub-figures.

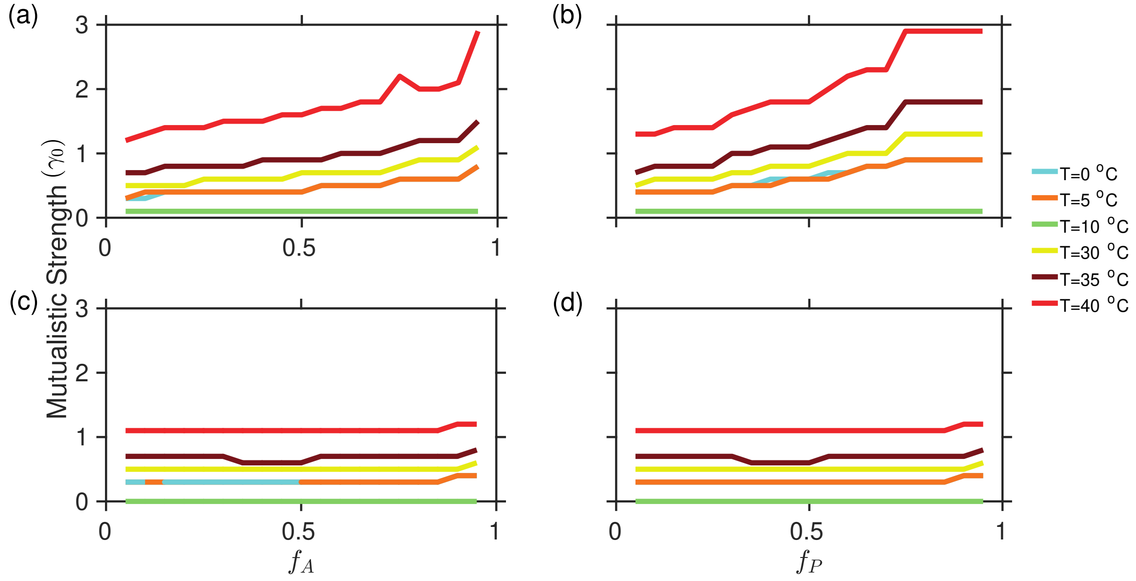


**Figure S1.13. Effects of pollinator and plant loss for varied temperature**: A fraction of (a) generalist pollinators ($f_A$), and (b) generalist plants ($f_P$) are removed in decreasing order of their degree at temperatures ranging over the interval $0 - 40^oC$. A fraction of (c) specialist pollinators ($f_A$), and (d) specialist plants ($f_P$) are removed in increasing order of their degree from a community. Generalist plant loss has a more significant effect on the collapse of the plant-pollinator community. A large difference is observed in tipping at higher temperatures for increase in $f_P$ and $f_A$ on removal of generalists populations in compared to specialists. Parameter values same as Fig. 1 (main paper)

**Table S1.1. Networks and their properties**: We present nestedness, modularity and connectance value corresponding to 16 different networks. $S_A$ and $S_P$ stand for the number of pollinators and plants, respectively.

| Sl. No. | Network's Name | $S_A$ | $S_P$ | Nestedness | Modularity | Connectance |
|---|---|---|---|---|---|---|
| 1. | $M\_PL\_061\_33$ | 6 | 2 | 0 | 0.2778 | 0.50 |
| 2. | $M\_PL\_056$ | 365 | 91 | 0.07 | 0.5267 | 0.0262 |
| 3. | $M\_PL\_030$ | 53 | 28 | 0.116 | 0.5841 | 0.0735 |
| 4. | $M\_PL\_002$ | 64 | 43 | 0.1536 | 0.5113 | 0.0712 |
| 5. | $M\_PL\_072\_01$ | 67 | 39 | 0.2016 | 0.47 | 0.0719 |
| 6. | $M\_PL\_061\_14$ | 11 | 6 | 0.2500 | 0.4599 | 0.2727 |
| 7. | $M\_PL\_051$ | 90 | 14 | 0.3001 | 0.4897 | 0.1302 |
| 8. | $M\_PL\_061\_05$ | 22 | 12 | 0.3539 | 0.4245 | 0.1932 |
| 9. | $M\_PL\_061\_24$ | 18 | 11 | 0.4028 | 0.4565 | 0.2323 |
| 10. | $M\_PL\_061\_07$ | 23 | 9 | 0.4589 | 0.29898 | 0.299 |
| 11. | $M\_PL\_064$ | 8 | 14 | 0.5049 | 0.3789 | 0.2857 |
| 12. | $M\_PL\_032$ | 33 | 7 | 0.5666 | 0.3477 | 0.2814 |
| 13. | $M\_PL\_066$ | 5 | 31 | 0.6299 | 0.2038 | 0.4710 |
| 14. | $M\_PL\_063$ | 9 | 55 | 0.6615 | 0.2903 | 0.2483 |
| 15. | $M\_PL\_069\_03$ | 4 | 7 | 0.7593 | 0.2000 | 0.5357 |
| 16. | $M\_PL\_059$ | 13 | 13 | 0.8493 | 0.2115 | 0.4207 |

**Table S1.2. Taxonomy details of an exemplary plant pollinator network**: Details corresponding to species constituting $M\_PL\_006$, species degree, and their count in other mutualistic networks.

| Specie | Kingdom | Degree | Networks presence |
|---|---|---|---|
| Animalsdela rufimatrella | Animals | 1 | 1 |
| Animalsndrena haemorrhoa | Animals | 3 | 3 |
| Animalsndrena pubescens | Animals | 2 | 2 |
| Animalsndrena wilkella | Animals | 5 | 3 |
| Animalsnosymia sp1 M_$PlantsL$_006 | Animals | 1 | 1 |
| Animalsnthomyidae sp1 M_$PlantsL$_006 | Animals | 3 | 1 |
| Animalsutographa sp1 M_$PlantsL$_006 | Animals | 1 | 1 |
| Bombus hortorum | Animals | 5 | 7 |
| Bombus lapidarius | Animals | 1 | 9 |
| Bombus pascuorum | Animals | 10 | 10 |
| Bombus pratorum | Animals | 3 | 7 |
| Bombus terrestris/lucorum | Animals | 10 | 2 |
| Cheilosia albitarsis | Animals | 2 | 4 |
| Chloromyia formosa | Animals | 1 | 2 |
| Chrysogaster sp1 M_$PlantsL$_006 | Animals | 1 | 1 |
| Empis livida | Animals | 1 | 2 |
| Eriothrix rufomaculatus | Animals | 1 | 3 |
| Eristalinus sepulcharius | Animals | 4 | 1 |
| Eristalis intricarius | Animals | 4 | 2 |
| Eristalis nemorum | Animals | 2 | 2 |
| Eristalis pertinax | Animals | 1 | 5 |
| Eristalis tenax | Animals | 2 | 22 |
| Helophilus sp1 M_$PlantsL$_006 | Animals | 8 | 1 |
| Helophilus trivittatus | Animals | 1 | 4 |
| Lasioglossum calceatum | Animals | 1 | 3 |
| Lasioglossum leucozonium | Animals | 1 | 3 |
| Lasioglossum villosulum | Animals | 1 | 6 |
| Lasiommata megera | Animals | 3 | 1 |
| Lejogaster splendida | Animals | 1 | 1 |
| Lucillia sp1 M_$PlantsL$_006 | Animals | 2 | 1 |
| Lycaena phlaeas | Animals | 1 | 8 |
| Malachius viridus | Animals | 1 | 1 |
| Maniola jurtina | Animals | 7 | 10 |
| Megachile willughbiella | Animals | 1 | 2 |
| Melanostoma sp1 M_$PlantsL$_006 | Animals | 1 | 1 |
| Meliscaeva sp1 M_$PlantsL$_006 | Animals | 1 | 1 |

| Specie | Kingdom | Degree | Networks presence |
|---|---|---|---|
| Micromoth sp1 M_$PlantsL$_006 | Animals | 1 | 1 |
| Nemotelus pantherinus | Animals | 1 | 1 |
| Neoascia tenur | Animals | 1 | 2 |
| Ochlodes venata | Animals | 2 | 2 |
| Odontomyia tigrina | Animals | 2 | 1 |
| Odontomyia viridula | Animals | 2 | 1 |
| Oligia sp1 M_$PlantsL$_006 | Animals | 1 | 1 |
| Plantsarhelophilus sp1 M_$PlantsL$_006 | Animals | 1 | 1 |
| Plantshaonia incarna | Animals | 8 | 1 |
| Plantslatycheirus sp1 M_$PlantsL$_006 | Animals | 2 | 1 |
| Plantsollenia sp1 M_$PlantsL$_006 | Animals | 2 | 1 |
| Plantsolyommatus icarus | Animals | 2 | 9 |
| Plantssithyrus vest | Animals | 2 | 2 |
| Plantsyronia tithonus | Animals | 1 | 2 |
| Rhagio tringarius | Animals | 1 | 1 |
| Rhagonycha fulva | Animals | 1 | 4 |
| Sarcophagus sp1 M_$PlantsL$_006 | Animals | 1 | 1 |
| Scathophaga stercoraria | Animals | 2 | 2 |
| Sphaerophoria sp1 M_$PlantsL$_006 | Animals | 3 | 1 |
| Syritta pipiens | Animals | 2 | 12 |
| Syrphus sp1 M_$PlantsL$_006 | Animals | 3 | 1 |
| Tetanocera ferruginea | Animals | 1 | 1 |
| Thymelicus sp1 M_$PlantsL$_006 | Animals | 3 | 1 |
| Tropidia scitta | Animals | 3 | 2 |
| Zygaena filipendulae | Animals | 4 | 4 |
| Animalschillea millefolium | Plants | 6 | 7 |
| Centaurea nigra | Plants | 3 | 3 |
| Cirsium arvense | Plants | 1 | 7 |
| Hypochoeris radicata | Plants | 15 | 6 |
| Lathyrus pratensis | Plants | 6 | 5 |
| Leucanthemum vulgare | Plants | 49 | 3 |
| Lotus corniculatus | Plants | 24 | 4 |
| Lychnis flos-cuculi | Plants | 1 | 4 |
| Plantslantago lanceolata | Plants | 2 | 1 |
| Plantsrunella vulgaris | Plants | 4 | 7 |
| Ranunculus acris | Plants | 12 | 3 |
| Taraxacum officinale | Plants | 1 | 6 |
| Trifolium dubium | Plants | 6 | 4 |
| Trifolium pratense | Plants | 2 | 5 |
| Trifolium repens | Plants | 1 | 11 |
| Vicia cracca | Plants | 9 | 2 |
| Vicia sativa | Plants | 4 | 1 |

### S2: Dimension reduction of the network model

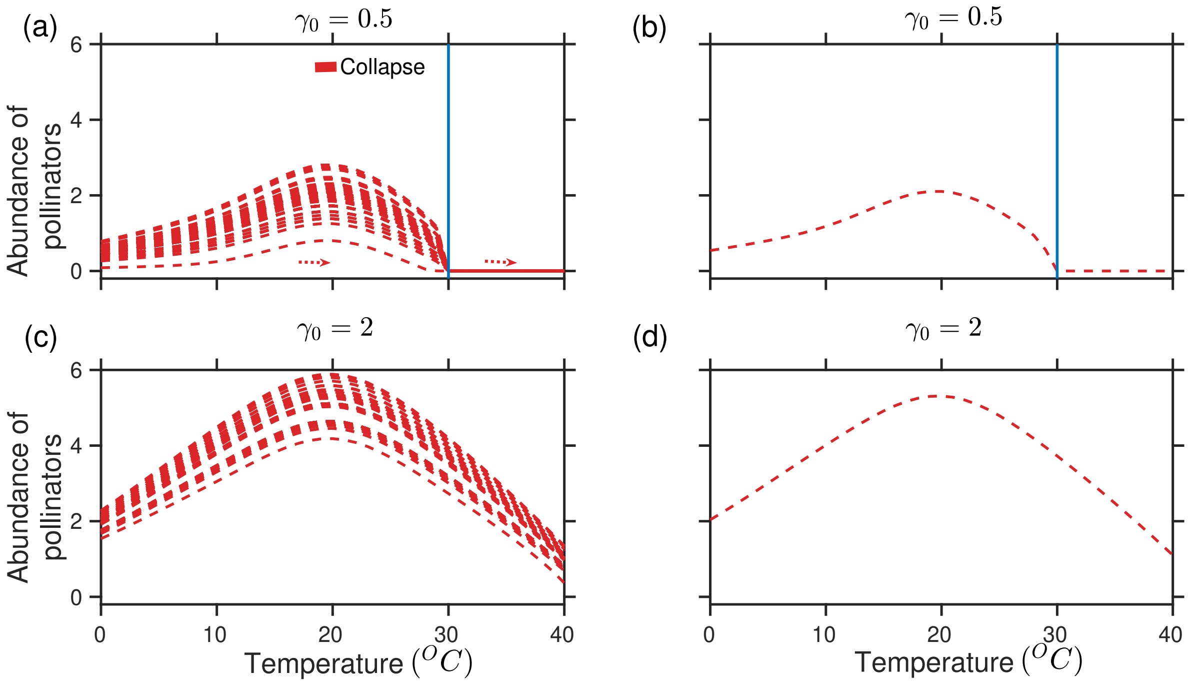


**Figure S2.1. Validation of reduced model dynamics**: Pollinator abundance plotted for increasing temperature in the range $0 - 40^0C$ (a)-(c) higher dimensional model (b)-(d) dimension reduced model. As observed dynamics close to tipping are well depicted by the dimension reduced model. The blue solid line represent the occurrence of a critical transition.

The average abundance of plants and pollinators obtained from Eq. 1 (main draft) can be expressed by: $\alpha_i^A(T)A_i \approx \alpha(T)A_e$, $\alpha_i^P(T)P_i \approx \alpha(T)P_e$, where $P_e$ and $A_e$ are the average abundance of plants and pollinators. When mutualistic partners are absent, species do not out-compete each other (van Nes and Scheffer 2004). Here we assume the intraspecific competition to be stronger than the interspecific competition, hence we consider:

$$\beta_{ii}^A >> \beta_{ij}^A \; ; \; \beta_{ii}^P >> \beta_{ij}^P \; .$$

Here we have neglected interspecific competition for simplicity to obtain $\beta_{ii}^A = \beta_{ii}^P = \beta = 1$ and $\beta_{ij}^A = \beta_{ij}^P = 0$. Therefore we have

$$\sum_{j=1}^{S_A} \beta_{ij}^A A_i A_j \approx \beta_{ii}^A A_e^2 = \beta A_e^2; \;\; \sum_{j=1}^{S_P} \beta_{ij}^P P_i P_j \approx \beta_{ii}^p P_e^2 = \beta P_e^2.$$

Incorporating interspecific competition term, the species competition can be represented as:

$$\sum_{j=1}^{S_A} \beta_{ij}^A A_i A_j \cong \frac{\sum_{i=1}^{S_A} \sum_{j=1}^{S_A} \beta_{ij}^A}{\sum_{i=1}^{S_A} 1} A_e^2 = \beta_A A_e^2,$$

$$\sum_{j=1}^{S_P} \beta_{ij}^P P_i P_j \cong \frac{\sum_{i=1}^{S_P} \sum_{j=1}^{S_P} \beta_{ij}^P}{\sum_{i=1}^{S_P} 1} P_e^2 = \beta_P P_e^2.$$

To calculate the actual mutualistic interaction in the networks, we have averaged mutualistic strength corresponding to each species in the following manner:

$$\sum_{j=1}^{S_P} \gamma_{ij}^A P_j = \sum_{j=1}^{S_P} \frac{\gamma_0}{d_{A_i}^{\delta}} \epsilon_{ij} P_j \cong \gamma_0 d_{A_i}^{1-\delta} P_e; \;\; d_{A_i} = \sum_{i=1}^{S_P} \epsilon_{ij},$$

$$\sum_{j=1}^{S_A} \gamma_{ij}^P A_j = \sum_{j=1}^{S_A} \frac{\gamma_0}{d_{P_i}^{\delta}} \epsilon_{ij} A_j \cong \gamma_0 d_{P_i}^{1-\delta} A_e; \;\; d_{P_i} = \sum_{i=1}^{S_A} \epsilon_{ij}.$$

There exist a variety of averaging method (Jiang et al 2018). Here we apply mainly three types of averaging methods namely unweighted, degree weighted, and eigenvector weighted methods to calculate average mutualistic strength in the system (Saavedra et al 2013). The dynamics is well depicted by the reduced model (Fig. S2.1) for each of the averaging method. The unweighted method is as follows:

$$\langle \gamma_A \rangle = \frac{\sum_{i=1}^{S_A} \gamma_0 d_{A_i}^{1-\delta}}{\sum_{i=1}^{S_A} 1},$$

$$\langle \gamma_P \rangle = \frac{\sum_{i=1}^{S_P} \gamma_0 d_{P_i}^{1-\delta}}{\sum_{i=1}^{S_P} 1}.$$

For the degree weighted method, we have

$$\langle \gamma_A \rangle = \frac{\sum_{i=1}^{S_A} \gamma_0 d_{A_i}^{1-\delta} \times d_{A_i}}{\sum_{i=1}^{S_A} d_{A_i}},$$

$$\langle \gamma_P \rangle = \frac{\sum_{i=1}^{S_P} \gamma_0 d_{P_i}^{1-\delta} \times d_{P_i}}{\sum_{i=1}^{S_P} d_{P_i}}.$$

The number of links for species associated with $A_i$ and $P_i$ are $d_{A_i}$ and $d_{P_i}$, respectively. For the eigenvector weighted method, we calculate the averaged quantities for plants and pollinators based on the eigenvector associated with the largest eigenvalue of the respective projection networks. Let $M_P$ and $M_A$ be the projection matrix of the plants and pollinators respectively. Then we have

$$M_P = M^T \times M; V_P = \text{eigenvector}(M_P),$$

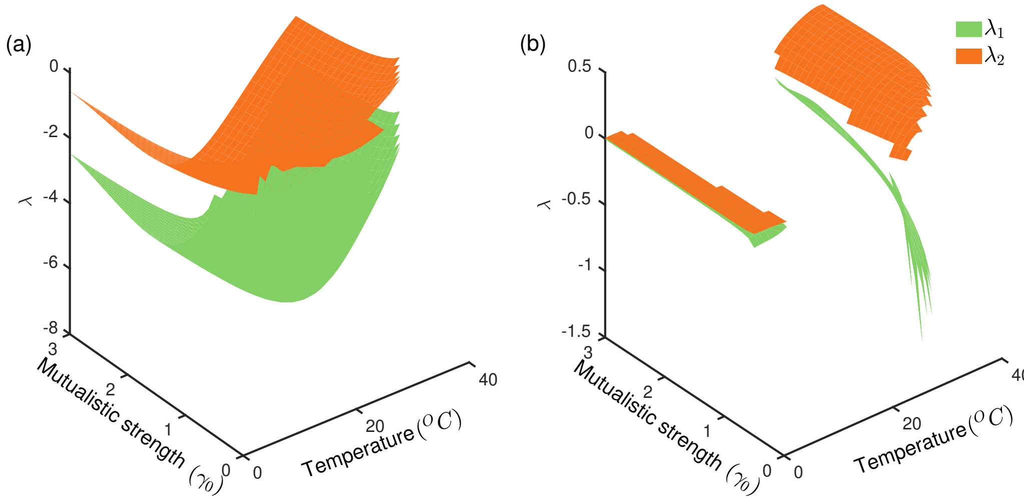


**Figure S2.2. Stability of the steady states of the reduced model versus the Mutualistic strength $\gamma_0$ and Temperature**: Green and orange surfaces represent the two eigenvalues of the Jacobian matrix evaluated at the steady state of the reduced model constructed from the empirical network. The figure represents the eigenvalue of SSS and USS versus temperature and mutualistic strength.

$$M_A = M \times M^T; V_A = \text{eigenvector}(M_A),$$

where $M$ is the $m \times n$ matrix characterizing the original bipartite network with $m$ and $n$ being the number of pollinators and plants, respectively. $V_A$ and $V_P$ are the components of the eigenvector associated with the largest eigenvalue of $M_A$ and $M_P$, respectively. The eigenvector weighted method is as follows:

$$\langle \gamma_A \rangle = \frac{\sum_{i=1}^{S_A} \gamma_0 d_{A_i}^{1-\delta} \times V_{A_i}}{\sum_{i=1}^{S_A} V_{A_i}},$$

$$\langle \gamma_P \rangle = \frac{\sum_{i=1}^{S_P} \gamma_0 d_{P_i}^{1-\delta} \times V_{P_i}}{\sum_{i=1}^{S_P} V_{P_i}},$$

where $V_{A_i}$ and $V_{P_i}$ are the $i^{th}$ component of $V_A$ and $V_P$, respectively.

The reduced model can be written as:

$$\frac{dA_e}{dt} = (\alpha(T) - k(T))A_e - \beta A_e^2 + \frac{\langle \gamma_A \rangle P_e}{1 + h(T)\langle \gamma_A \rangle P_e} A_e + \mu, \tag{1a}$$

$$\frac{dP_e}{dt} = \alpha(T)P_e - \beta P_e^2 + \frac{\langle \gamma_P \rangle A_e}{1 + h(T)\langle \gamma_P \rangle A_e} P_e + \mu, \tag{1b}$$

where $P_e$ and $A_e$ are the average plant and pollinator densities, respectively. $\alpha$, $\beta$, $h$, $\langle \gamma_A \rangle$, $\langle \gamma_P \rangle$, and $\mu$ denote the mean of the corresponding parameters in Eq. 1 (main draft), and have the same interpretation. $\langle\gamma_P\rangle$, and $\langle\gamma_A\rangle$ are calculated using an averaging method (Jiang et al 2018) (for further details see *SI Appendix,* Section S2). Here, we use the eigenvector weighting method for finding $\langle\gamma_P\rangle$ and $\langle\gamma_A\rangle$ which also fits the $n$-th dimensional model. Equating right hand side of Eq. (1), we have calculated the steady state solution for a particular temperature. The steady state solution is given by:

$$A_s \approx \frac{1}{\beta}\left[\alpha(T) - k(T) + \frac{\langle\gamma_A\rangle P_s}{1 + h(T)\langle\gamma_A\rangle P_s}\right], \text{ and}$$

$$P_s \approx \frac{1}{\beta}\left[\alpha(T) + \frac{\langle\gamma_P\rangle A_s}{1 + h(T)\langle\gamma_P\rangle A_s}\right].$$

The equation of $A_s$ becomes: $c_1(T)A_s^2 + c_2(T)A_s + c_3(T) = 0$, where

$$\begin{aligned}
c_1(T) &= -\left(\beta^2 h(T)\langle\gamma_P\rangle + \beta h(T)\langle\gamma_A\rangle\langle\gamma_P\rangle + \beta h^2(T)\alpha(T)\langle\gamma_A\rangle\langle\gamma_P\rangle\right),\\
c_2(T) &= \beta^2 - \beta h(T)\alpha(T)\left(\langle\gamma_A\rangle - \langle\gamma_P\rangle\right) + \langle\gamma_A\rangle\langle\gamma_P\rangle\left(1 + h(T)\alpha(T)\right)^2\\
&\quad -k(T)\left(\beta h(T)\langle\gamma_P\rangle + h(T)\langle\gamma_A\rangle\langle\gamma_P\rangle + h^2(T)\alpha(T)\langle\gamma_A\rangle\langle\gamma_P\rangle\right), \text{ and}\\
c_3(T) &= \alpha(T)\beta + \alpha(T)\langle\gamma_A\rangle + h(T)\alpha^2(T)\langle\gamma_A\rangle - k(T)\left(\beta + h(T)\alpha(T)\langle\gamma_A\rangle\right).
\end{aligned}$$

Averaged pollinator abundance for unstable steady state ($USS$) and stable steady state ($SSS$) are given below:

$$A_{USS} = \frac{-c_2(T) + \sqrt{c_2^2(T) - 4c_1(T)c_3(T)}}{2c_1(T)}, \text{ and } A_{SSS} = \frac{-c_2(T) - \sqrt{c_2^2(T) - 4c_1(T)c_3(T)}}{2c_1(T)}$$

Averaged plant abundance for $USS$ and $SSS$ are as follows:

$$P_{USS} = \alpha(T) + \frac{\langle\gamma_P\rangle A_{USS}}{1 + h(T)\langle\gamma_P\rangle A_{USS}}, \text{ and } P_{SSS} = \alpha(T) + \frac{\langle\gamma_P\rangle A_{SSS}}{1 + h(T)\langle\gamma_P\rangle A_{SSS}}.$$

Further, we have studied the SSS and USS using the eigenvalue of the corresponding Jacobian matrix for the reduced model at the steady states (see Fig. S2.2).
